# Defined microenvironments trigger *in vitro* gastrulation in human pluripotent stem cells

**DOI:** 10.1101/2021.10.28.466327

**Authors:** Pallavi Srivastava, Sara Romanazzo, Jake Ireland, Stephanie Nemec, Thomas G. Molley, Pavithra Jayathilaka, Elvis Pandzic, Avani Yeola, Vashe Chandrakanthan, John Pimanda, Kristopher Kilian

## Abstract

Embryogenesis is orchestrated through local morphogen gradients and endometrial constraints that give rise to the three germ layers in a well-defined assembly. *In vitro* models of embryogenesis have been demonstrated by treating pluripotent stem cells in adherent or suspension culture with soluble morphogens and small molecules, which leads to tri-lineage differentiation. However, treatment with exogenous agents override the subtle spatiotemporal changes observed *in vivo* that ultimately underly the human body plan. Here we demonstrate how microconfinement of pluripotent stem cells on hydrogel substrates catalyses gastrulation-like events without the need for supplements. Within six hours of initial seeding, cells at the boundary show elevated cytoskeletal tension and yes-associated protein (YAP) activity, which leads to changes in cell and nuclear morphology, epithelial to mesenchymal transition, and emergence of defined patterns of primitive streak containing SRY-Box Transcription Factor 17 (SOX17)^+^ T/BRACHYURY^+^ cells. Immunofluorescence staining, transcript analysis, and the use of pharmacological modulators reveal a role for mechanotransduction-coupled non-canonical wingless-type (WNT) signalling in promoting epithelial to mesenchymal transition and multilayered organization within the colonies. These microscale gastruloids were removed from the substrate and encapsulated in 3D hydrogels, where biomaterials properties correspond to maintenance and spatial positioning of the primitive streak. Together, this approach demonstrates how materials alone can nurture embryonic gastrulation, thereby providing an *in vitro* model of early development.

## Introduction

One of the most important events in developmental biology is gastrulation, where single layered pluripotent epiblast cells go through a series of carefully regulated cell fate decisions to form the progenitors of the three germ layers ^1^. Previous studies in mouse have revealed that the posterior side of the embryo initiates a region of cells undergoing epithelial-to-mesenchymal transition (EMT) to form the primitive streak, and delaminate from the epiblast surface after gaining mesenchymal motility^2^. The cells in this primitive streak region are positive for the mesodermal marker T/BRACHYURY, and as they ingress inwards, they segregate towards endoderm progenitors with endodermal marker SOX17 as the streak extends. These mes-endodermal cells further proceed to ingress through the primitive streak into the gastrulating embryo^3^. The dramatic cellular identity changes *in vivo* are attributed to an interplay of the TGFβ, WNT and FGF signalling pathways with their respective antagonists^4^ and their order of activation and patterns of gene regulation have been well studied^5^.

Recently, there has been an effort to understand developmental signals in the context of biophysical forces during gastrulation; specifically, the local mechanical forces and geometric constraints created by surrounding extra-embryonic tissues during embryo implantation into the uterine lining^6^. Biophysical regulation of the gastrulation process is difficult to study due to ethical and physiological limitations with handling human embryos after the appearance of the primitive streak, approximately 14 days after fertilisation^7^. The anatomy of embryonic development contains several phases where extra-embryonic pressure and the mechanics of the surrounding tissue will impose constraints on morphogenesis. However, current opinion is mixed regarding the importance of the biophysical microenvironment in directing early stages of embryogenesis. Therefore, *in vitro* approaches to culture or encapsulate pluripotent stem cells or epiblast stem cells, using hydrogels, microcarriers, scaffolds and other biomaterials have been used to model gastrulation *in vitro* towards understanding the role biophysical factors play during embryogenesis^8, 9^.

I*n vitro* models for gastrulation using microengineering have allowed the effect of geometric confinement during tri-lineage differentiation to be probed^10, 11^. After bone morphogenic protein-4 (BMP4) stimulation for 48 hours, pluripotent stem cells (PSCs) undergo spatial patterning in response to confinement, which is reminiscent of *in vivo* processes. Most *in vitro* studies use rigid glass surfaces (∼3-4 GPa), and exogenous supplements of soluble morphogens to trigger differentiation. In contrast, the signalling gradients *in vivo* are dynamic, involving local gradients of paracrine and autocrine signals to drive morphogenesis in a confined microenvironment with variable viscoelastic properties^12^. Pre-streak formation events involve actin-controlled oriented cell movements and deformation on the posterior epiblast, which guide dynamic local gradients of signalling to coordinate primitive streak progression.^13, 14^ Recently, Weaver and colleagues reported the appearance of multicellular T/BRACHYURY^+^ “gastrulation-like” mesodermal nodes at the colony edges on soft patterned substrates following BMP4 driven differentiation, highlighting the interplay between mechanics and morphogens^15^. These *in vitro* gastruloids show similarities in gene expression and cellular organization with several hallmark features of gastrulation – mes-endodermal Identity (along with comparable downregulation of pluripotent identity), evidence of EMT, and collective cell movement after loss of pluripotency^15^.

In this article we demonstrate how human pluripotent stem cells spontaneously differentiate into SOX17^+^ T/BRACHYURY^+^ gastruloid-like structures within 2 days when microconfined on precise matrix-conjugated deformable substrates, without the need for exogenous supplements. The degree of lineage specification corresponds to matrix biophysical conditions, with a decline in pluripotency coinciding with EMT and mes-endodermal gene activation. Treatment with BMP4 enhances this transformation, resulting in the appearance of textured wave-like regions of multilayered mes-endodermal SOX17^+^ and T/BRACHYURY^+^ structures within the micropatterned colonies. Gene expression analysis coupled with pathway inhibition studies indicates that mechanotransduction triggers gastrulation in confined colonies through non-canonical wingless-type (WNT) signalling. Release from confinement and encapsulation in tailored hydrogels results in spatial patterning of differentiation, thereby providing a “materials-centric” method of forming gastrulation mimics to model embryogenesis.

## Results

### Protein-conjugated hydrogels facilitate human induced pluripotent stem cell (iPSCs) culture

To assess the behaviour of hiPSCs on compliant substrates, we chose to work with polyacrylamide (PA) hydrogels with tuneable stiffness that can be further modified using soft lithography to define regions of adhesivity^16^. Briefly, the concentration of acrylamide and bis-acrylamide was varied to create PA gel solutions for stiffness of 1, 10 and 100 kPa (10-fold increase spanning physiological tissue) and then polymerised on chemically modified glass coverslips. The surface of the PA was treated with hydrazine hydrate and then imprinted with an oxidized protein using soft lithography to form the covalent Schiff base using patterned or flat polydimethylsiloxane stamps to mediate cell adhesion on an otherwise inert PA surface. The same stamps were used to assist with physical adsorption of the protein on a clean glass surface to define protein islands. For this study, we had four test groups: protein coated glass, protein patterned glass, protein coated hydrogels and protein patterned hydrogels (hydrogels formulated at 1, 10 and 100 kPa). These groups assessed the response of iPSCs towards confinement alone, stiffness alone and geometry combined with stiffness, all compared to bare glass controls. To screen optimal proteins for iPSC attachment and proliferation, we assayed common matrix proteins involved in *in vitro* stem cell culture: EHS Laminin, Laminin-521, Collagen, Fibronectin, hESC qualified Matrigel and rh-Vitronectin, deposited on substates via soft lithography (Figure S1A). These trials revealed rh-Vitronectin to be the most compatible protein which readily facilitated iPSC attachment and proliferation (Figure 1A), as compared to all other tested proteins where high cell death or little to no cell attachment was observed. Since the microcontact patterning process of the PA hydrogels requires an oxidation of the printing protein using sodium periodate at room temperature, rh-Vitronectin, which is generally stable at room temperature proved to be a better alternative to matrigel, which was unstable in supporting patterned iPSC adhesion under these conditions.

**Figure 1:**
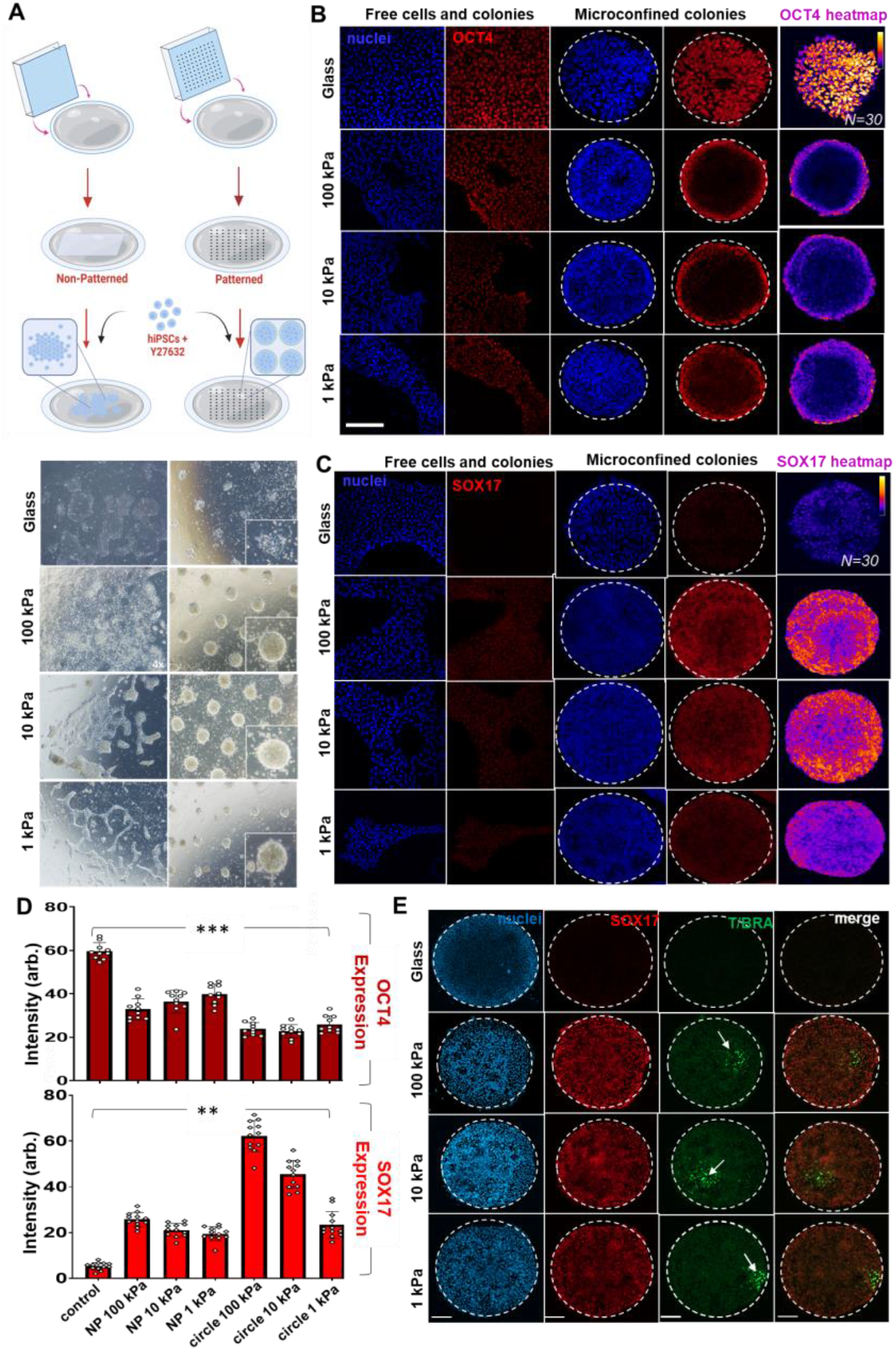
Substrate stiffness directs SOX17^+^ T/BRACHYURY^+^ mes-endodermal population. **A)** Schematic of PA hydrogel substrate preparation and cell seeding procedure, brightfield images from non-pattered and patterned 250µM colonies at 48 hours. All images acquired at 4x objective. **B)** hiPSCs seeded non-patterned and patterned glass and hydrogel substrates immunostained for Oct4. Right –immunofluorescence heatmaps of Oct4 expression. **C)** hiPSCs seeded non-patterned and patterned glass and hydrogel substrates stained for Sox17. Right –immunofluorescence heatmaps of Sox17 expression. **D)** Comparison of expression intensity for OCT4 and SOX17 across all conditions (N=8). *** = p<0.001,**=p<0.01 (Ordinary One-way ANOVA) **E)** Immunofluorescence images of endoderm marker SOX17 and mesoderm marker T/Bra in cell populations on 500µM circular colonies. Scale Bars: 100um

Having identified vitronectin as a suitable protein for hiPSC culture, we next asked whether these hydrogel matrices would maintain the pluripotent phenotype. The hiPSCs on glass showed positivity for pluripotency markers (OCT4 and NANOG) and were simultaneously found to be negative for germ layer lineage markers (Figure S1B). hiPSCs demonstrated healthy attachment on the non-patterned and patterned PA substrates, with adherent colonies initiating multilayered growth after 48 hours.

### Substrate properties alone guide human pluripotent stem cell lineage specification towards mes-endodermal identity

Having established optimal conditions for iPSC culture on hydrogels, we next sought to assess the expression of molecular markers of pluripotency and tri-lineage differentiation. For geometric confinement, 250µM and 500µM diameter circles were tested. The hiPSCs were dissociated to single cells and seeded at a uniform cell density while being supplemented with Y-27632 and allowed to grow for 48h before fixation. Optimal cell density was selected based on conditions that foster near confluence on day one.

To determine the pluripotency status and lineage identity of hiPSCs on these different surfaces, we immunostained the cell populations with the pluripotency marker OCT4 (POU5f1), as well as definitive endoderm marker – SOX17^17^ and mesoderm/primitive streak marker – T/BRACHYURY^5^. We observed that the cells cultured on glass controls and glass patterns, maintained pluripotency with uniform expression of OCT4 (Figure 1B). However, hiPSCs cultured on non-patterned hydrogels across each stiffness condition demonstrated decreased expression of OCT4. Microconfined cells across all three stiffnesses demonstrated OCT4 expression restricted to the colony edges, with complete loss of signal in the centre (Figure 1B,D). This appearance of an OCT4 annulus in a confined colony was observed previously during cardiac differentiation on patterned glass using CHIR99021 induction,^18^ which was postulated to be a Wnt signaling-mechanics relationship. Expression of the endoderm marker SOX17 was negligible in colonies on glass, with evidence for modest expression in colonies on non-patterned hydrogels, indicating a potential role for substrate mechanics in regulating endoderm specification. In contrast, there was a striking upregulation of SOX17 in hydrogel microconfined conditions, with the highest expression observed in cells on 10 and 100 kPa, with decreased expression in cells confined on 1 kPa hydrogels (Figure 1C, D). Substrate softness has been shown to favour endodermal differentiation.^19, 20^ However, here we see that confining hiPSCs on softer substrates created a multilayered endodermal structure which contained a small mesodermal cell cluster within. The SOX17 expression in colonies confined to micropatterned hydrogels was observed as early as 12h after seeding (Figure S2), on both 250 µM and 500 µM diameter micropatterned hydrogels, with the majority of the cells co-expressing the endodermal marker FOXA2 (Figure S3). To rule out artefacts associated with reprogrammed cells, we cultured H9 human embryonic stem cells under the same conditions and observed a similar increased SOX17 expression on 10kPa hydrogels (Figure S4).

A distinct but transient expression of T/BRACHYURY was also identified in a small population of cells towards the colony centre, more commonly observed in the 10kPa condition, in about 40-50% of experimental replicates (Figure 1E), whereas distinct nuclear punctate were observed across all replicates of confined 100kPa and 10kPa hydrogel conditions, with localisation towards the colony centre (Figure S5). We also stained for the ectodermal marker SOX2, and neuroectoderm progenitor SOX1, with comparable negative staining across all confined hydrogel substrates. This shows that the colonies on PA hydrogels were mostly mes-endodermal in identity, and that primitive streak/gastruloid-like identity was instigated by the imposed biophysical microenvironment. To ensure reproducibility, all results were reproduced in at least 6 technical replicates with a minimum of 3 biological replicates for each experiment.

Previously we demonstrated how changes in perimeter curvature would influence the behaviour of microconfined cells, where geometry and stiffness both exerted an influence over cell phenotype in the context of cancer stemness.^21, 22^. To evaluate whether geometry would play a role in the observed differentiation with our microconfined hiPSC colonies, we cultured cells in shapes of the same area approximating a star, flower, square, and capital ‘I’—where positive and negative curvature and aspect ratio are varied—followed by immunostaining for germ layer markers. After 48 hours there were no significant differences between mes-endodermal marker expression across the shaped colonies, suggesting changes in geometry at the interface does not play a role in the observed gastrulation-like morphogenesis (Figure S6).

### Early adhesion stimulates YAP activity to coordinate epithelial-to-mesenchymal transition and differentiation

Having observed how hydrogel microconfinement alone will trigger mes-endodermal differentiation, we next sought to investigate how the surface directs this effect. Epiblast cells at the primitive streak region are widely reported to have undergone EMT, to gain their mesenchymal and mes-endodermal identity before they ingress in the gastrulating embryo^28^. We immunostained microconfined colonies on 10 kPa hydrogel and glass patterns for EMT molecular markers E-CAD, N-CAD and SNAIL, and OCT4 to gauge pluripotency. We selected the stiffness 10kPa for our experiments hereafter since we observed a more consistent spontaneous appearance of the SOX17^+^ colonies with the T/BRACHYURY^+^ clusters on 10 kPa as compared to 1 and 100 kPa. As before, we observed OCT4 expression being restricted towards the colony edges with decreased expression towards the centre (Figure 2 A,B). At the same time, there is a striking loss of E-CAD in the colony centres, suggesting an EMT prone region starting inwards from the periphery. In conjunction with loss of E-CAD, the population confined on hydrogels expresses uniform SNAIL, with N-CAD expression in the colony centre. This E-CAD to N-CAD switch is considered a prime indicator of cells undergoing EMT and is considered crucial for specification of the primitive streak and other embryogenesis events^29^. In comparison, E-CAD and OCT4 show uniform expression throughout the colonies on glass patterns, with no expression of SNAIL (Figure 2A). E-CAD expression on the cell membrane and cytoplasm was discontinuous in regions of the 250 µm colonies (Figure S7A). Moreover, E-cad expression was maintained in colonies across non-patterned surfaces, with only a few regions showing discontinuous or punctate E-CAD on the non-patterned hydrogels (Figure S7B).

**Figure 2:**
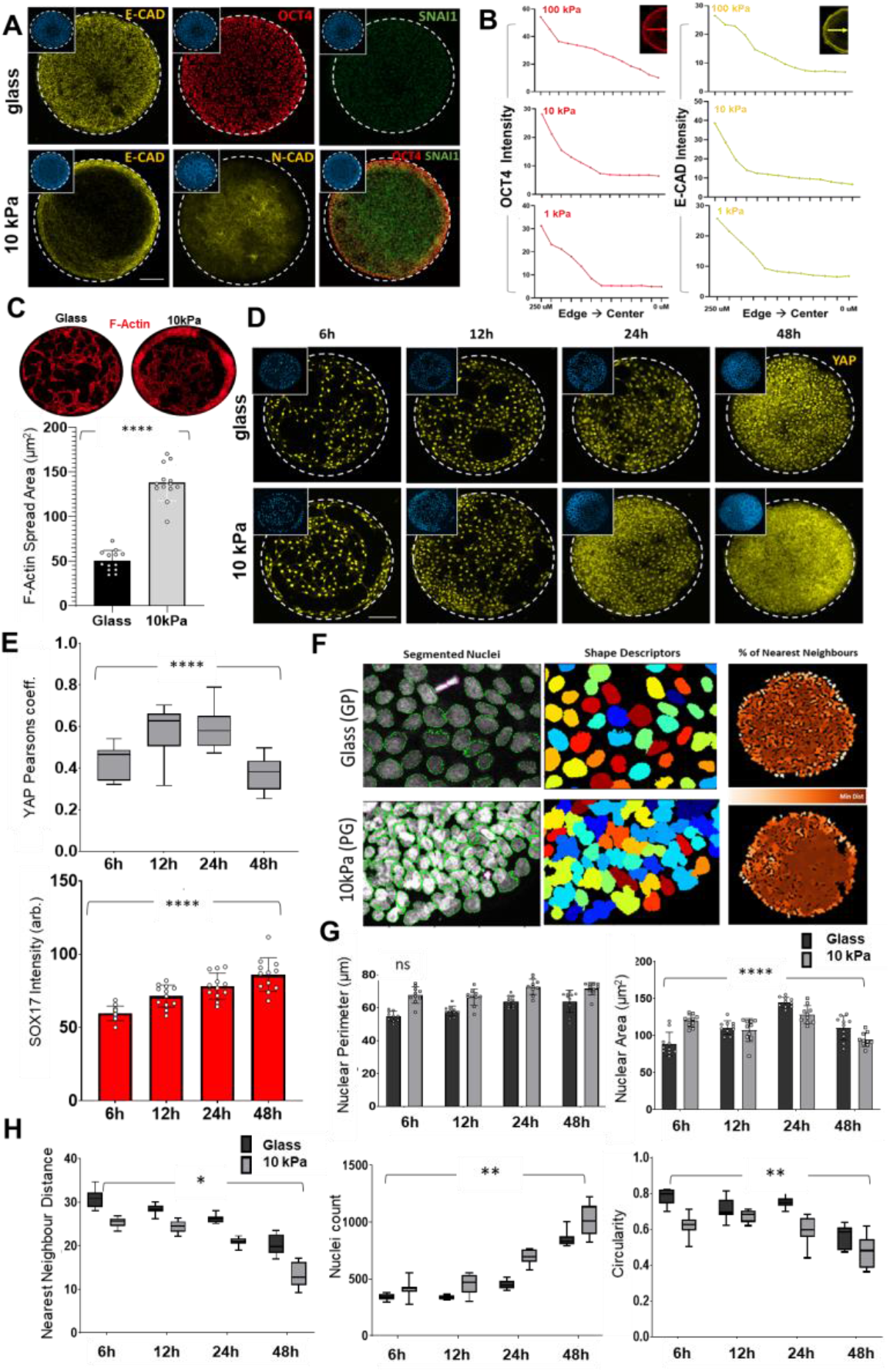
Early adhesion stimulates YAP activity to coordinate epithelial-to-mesenchymal transition and differentiation. **A)** Immunofluorescence images for E-CAD, N-CAD, SNAIL and OCT4 across colonies on glass and hydrogel patterns. **B)** Trace plots quantifying OCT4 and E-CAD expression on PA hydrogels from edge to centre. **C)** Actin spread area comparison for glass and 10kPa 500µM patterns. **** = p<0.0001 (N=13). **D)** Time course analysis of adhesion pattern and localisation of YAP on glass and 10kPa patterns. **E)** Quantification of YAP nuclear-cytoplasmic ratio (top) and corresponding SOX17 expression intensity (bottom) at four time points. **** = p<0.0001, ***=p<0.001, N=12 (Ordinary One-way ANOVA). **F)** Nuclei segmentation as primary objects, nuclei shape description, percentage depiction of nearest neighbours on glass and 10kPa 500µM patterns. *** = p<0.001,**=p<0.01(Two-way ANOVA) **G)** and **H)** Comparison of nuclei area and perimeter, distance between nearest neighbours, average cell count and averaged circularity of identified nuclei on glass and 10kPa 500µM patterns *** = p<0.001,**=p<0.01, *=p<0.05, ns=not significant (Two-way ANOVA). Scale bars = 100µM

We observed a region at the colony edge with a SNAIL^+^ OCT4^+^ population, which is also the region with highest SOX17 expression. We propose that this overlap denotes early stages in embryo development where SNAIL is reported to control EMT at the epiblast via downregulation of E-cadherin^30^. Similar SNAIL expression was also observed in the 250 µm colonies across all patterned hydrogels; however, there was no apparent spatial organization (Figure S7A). The E-CAD to N-CAD switch and concurrent SNAIL expression implicates EMT as the morphogenetic process that directs endodermal/mesodermal identity in the microconfined colonies on hydrogels.

Next, we analysed differences in cell adhesion and proliferation over time in our microconfined cultures by fixing and staining at 6, 12, 24, and 48 hours with the same initial seeding density. After initial seeding, cells encircle the border and adopt an elongated contractile morphology (Figure S2) with elevated F-actin and higher spread area compared to cells in the interior of the pattern or those constrained in glass patterns (Figure 2C). Over time, these contractile cells proliferate to fill the pattern which coincides with lessening of the observed cytoskeletal tension. Confined cells show significantly higher proliferation on the hydrogel patterns compared to glass, which at first glance is counterintuitive since soft matrices are known to limit cell proliferation. However, ROCK inhibition with Y27632 has been reported to promote cell proliferation on soft matrices and supress proliferation on glass through modulating actomyosin contractility^31^. This is also apparent through a nearest neighbours analysis (Figure 2 G,H). A core protein in sensing matrix stiffness, and a driver of pluripotency maintenance, is the yes-associated protein (YAP)^32^. YAP activity is involved in controlling the expression of genes associated with the anterior primitive streak^33^ and has been indicated in coordinating germ layer patterning during *in vitro* BMP4 driven gastrulation^34^. YAP expression is predominantly nuclear within the first 12 hours, followed by a decrease as the cells proliferate from the perimeter to the centre (Figure 3 D,E). Monitoring SOX17 expression at the same time showed that the transition of nuclear YAP to the cytoplasm corresponds with decreased cytoskeletal tension and increase SOX17 expression. During this time course we also measured changes in nuclear geometry, where variations in morphology has been demonstrated during EMT, differentiation and de-differentiation.^35, 36^ Cells confined on 10 kPa hydrogels show increased nuclear perimeter over time with decreased circularity compared to cells confined on glass (Figure 3 F,G,H). Overall, these results suggest that initial adhesion increases cytoskeletal tension at the boundary, directing YAP activity and stimulating EMT, followed by differentiation to a mes-endodermal population.

**Figure 3:**
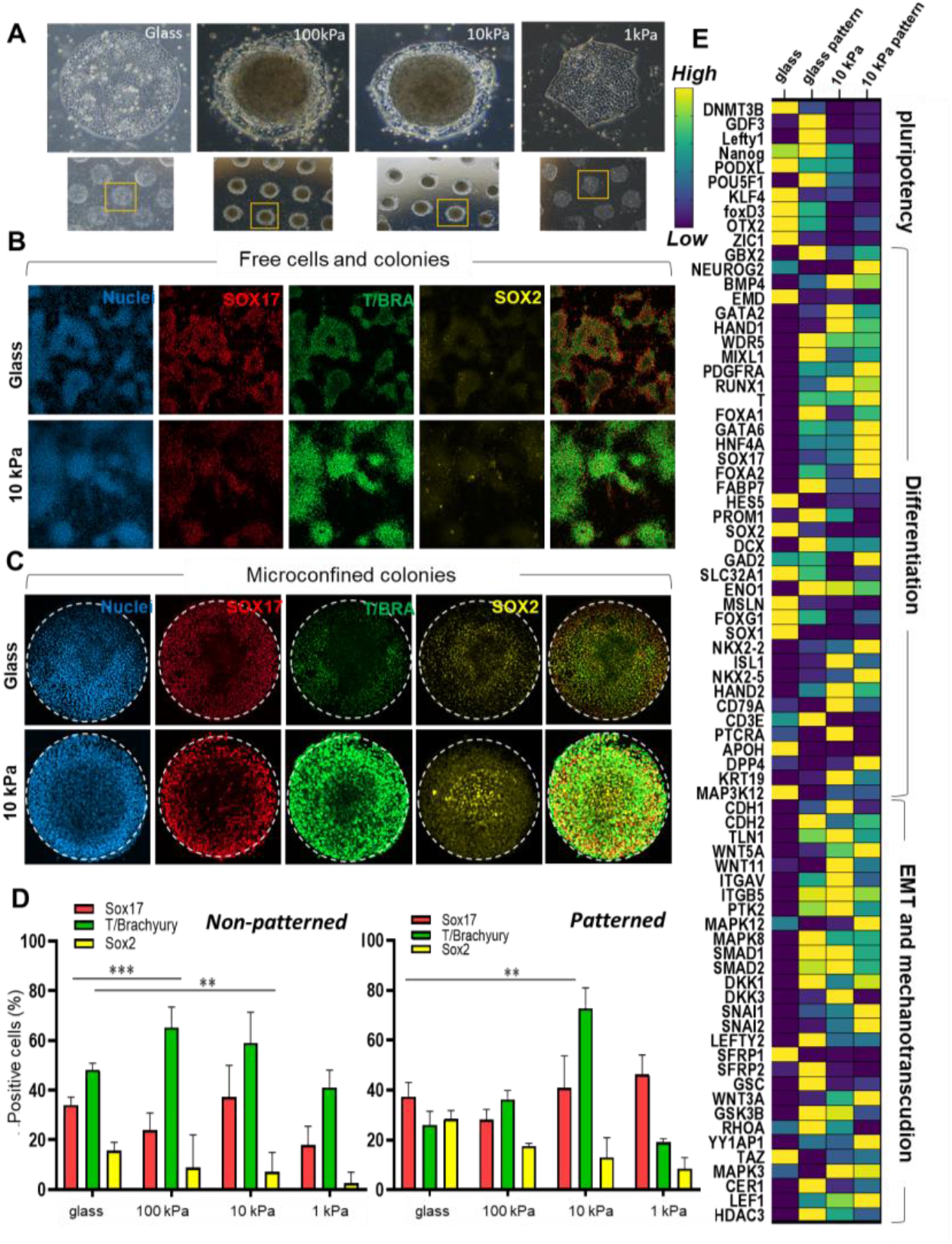
Priming pluripotent stem cell colonies augments BMP4 induced germ layer specification. **A)** Brightfield images of individual representative colonies on each substrate. **B)** BMP4 induction leads to gastruloid-like organized cell clusters on glass and hydrogel substrates (Scale bar – 250µM). **C)** Circular confinement centralizes the primitive streak-like cluster in the microconfined colonies (Scale bar – 100µM). **D)** Quantification of the percentage of positive cells for the three markers on non-patterned glass and hydrogels (top) and patterned glass and hydrogels (bottom) *** = p<0.001,**=p<0.01)(Two-way ANOVA) **E)** Heatmap of quantitative expression of genes associated with pluripotency, differentiation, EMT and mechanotransduction (N=2).

### Priming pluripotent stem cell colonies on hydrogels augments BMP4 induced germ layer specification

Previous work with micropatterned cultures used BMP4 as a soluble morphogen to initiate differentiation.^10, 11, 15^ We hypothesised that our cells on micropatterned hydrogels would be more susceptible to differentiation, or bias the population to different outcomes, after induction with BMP4 when compared to the standard condition of culture on glass. To test this, we seeded hiPSCs with BMP4 (50 ng/ml) induction to drive differentiation over 48 hours. In the brightfield images, the colonies on the glass surface show homogenous distribution with some multilayering at the edges. In contrast, the colonies on 10 and 100 kPa hydrogels showed distinct multilayered clustering towards the centre with monolayer organization at the border (Figure 3A). Cells confined on 1 kPa hydrogels did not show this multilayered characteristic. We immunostained all cultures for germ layer markers SOX17, T/BRACHYURY and SOX2. After BMP4 induction, the cells cultured on glass showed periodic patterns of clustered and spread cells with a wave-like morphology, where the edges of the clustered regions are rimmed with mes-endodermal SOX17^+^ T/BRACHYURY^+^ cells with slight regions of ectodermal SOX2^+^ (Figure 3B). This result demonstrated how BMP4 induction promotes outer endoderm with adjacent mesoderm and ectoderm, with concentric organization when constrained in glass micropatterns.^10^ The cells seeded on non-patterned hydrogels similarly showed wave-like multilayered texturing with increased density and increased mesoderm specification observed on the 10 kPa hydrogels. Cells cultured on the non-patterned 1 kPa hydrogel showed comparatively less attachment leading to smaller clusters of randomly placed mes-endodermal cells (Figure S8A). Strikingly, the concentric spatial patterning of germ layers in microconfined colonies on glass was also observed on 500 µM patterned hydrogels, but with a considerable enhancement leading to a large multilayered cluster in the colony centre (Figure 3C). These central clusters were dense with SOX17^+^ cells located towards the edge, intermingled with a population of T/BRACHYURY^+^ cells extending inward, with SOX2^+^ cells exclusively at the centre. We observed similar characteristics in cell colonies in the smaller 250 µM diameter patterns (Figure S8B); immunofluorescence quantitation revealed enhanced T/BRACHYURY+ mesoderm specification on the 10 kPa hydrogel micropatterns, confirming the previous observations of softer substrates favouring mesodermal differentiation in response to BMP4^15^ (Figure 3D).

Since EMT processes were observed in our cultures without BMP4 (Figure 2), we also immunostained these cultures for E-cadherin (E-CAD) which demonstrated a pronounced loss towards the colony edges on glass, with the reverse observed for colonies cultured in confinement on hydrogels (Figure S9A). These trends in E-CAD expression correspond directly with the expression of markers associated with differentiation. Another characteristic of differentiation is changes in cell and nuclear area. The area and perimeter of the differentiated cell nuclei was considerably lower in clustered cells compared to spread cells across all conditions (Figure S9B).

To support our immunofluorescence result, we selected our optimal hydrogel condition (10 kPa) compared to glass for quantitative PCR to assess differences in transcript expression on account of substrate stiffens, geometric confinement and both (Figure 3E). Consistent with the immunofluorescence results for microconfined colonies on hydrogels, we observed decreased expression of pluripotency genes with an increase in differentiation markers towards mes-endoderm lineages compared to colonies on glass—e.g., endoderm: SOX17 (x340), FOXA2 (x247) and GATA6 (x592), and mesoderm: T/BRACHYURY (x 43), RUNX1 (x33) and MIXL1 (x50). We also observed increased expression of EMT regulators N-CAD (CDH2; x4), SNAIL (SNAI1; x13); SNAI2 (x163), and downstream effectors of BMP signalling SMAD1/2 (x2) in the micropatterned colonies. Canonical WNT signalling member WNT3A (x28) and its inhibitor DKK1 (x72) were also considerably elevated on micropatterned hydrogels; both molecules playing central roles in controlling gastruloid pattern extension and meso-vs endo-differentiation^23^. Nodal antagonist CEREBRUS 1 (CER1) was considerably elevated in micropatterned colonies (x430), suggesting a role contributing to primitive streak-like formation in microconfined cultures.

In addition to pluripotency, differentiation and EMT markers, microconfinement on hydrogels led to increased expression of vitronectin binding integrin αV (x2) and β5 (x4) and associated downstream effectors of mechanotransduction, consistent with our observations of cytoskeletal tension and YAP activity guiding morphogenesis. Concurrently, we saw an increase in non-canonical WNT5A (x59) and WNT11 (x2), which are regulated by several mechanotransduction pathways including planar cell polarity which dictates patterning during embryogenesis in multiple species^24, 25^. Furthermore, there was 28-fold higher expression of the WNT inhibitor secreted frizzled related protein (SFRP1) in colonies on glass, suggesting attenuation of WNT signals is a central aspect of maintaining pluripotency. Previous reports demonstrate the importance of WNT in ES cell differentiation^26^ as well as the role of mechanics favouring mesoderm differentiation on compliant surfaces^15, 27^. Together these results demonstrate how microconfinement on hydrogels enhances EMT and WNT/Nodal activity which leads to increased differentiation upon treatment with BMP4.

### WNT signalling contributes to differentiation in spatially confined microenvironments

Pluripotent stem cells seeded on hydrogel micropatterns experience edge stress that increases cytoskeletal tension affecting YAP localisation, thereby initiating EMT and differentiation. While our gene expression analysis aligns with these mechanotransduction pathways guiding differentiation on hydrogels, there were also considerable changes in numerous paracrine signalling pathways involved in embryogenesis. The formation of primitive streak *in vivo* is guided by the activity of TGFβ/Nodal as well as WNT/β-Catenin pathways^15, 23^. Since these pathways have been shown to synergistically push the epiblast cells towards primitive streak while blocking ectoderm differentiation^37^, we next investigated the use of small molecule pharmacological disruptors to attenuate WNT and TGFβ signalling.

Canonical WNT signalling involves β-catenin activity and is associated with mesodermal differentiation,^38^ definitive endoderm progenitors,^40^ as well as EMT related gain of motility supporting primitive streak.^39^ Since our transcript analysis showed elevated WNT signalling, we supplemented our cultures with the canonical WNT inhibitor IWP2. Treatment with IWP2 leads to complete loss of T/BRACHYURY expression in colonies on the 10kPa hydrogel. However, the average SOX17 expression remains unchanged compared to untreated (Figure 4A). This result suggests that canonical WNT signalling is critical for mesoderm but not endoderm differentiation. Our transcript analysis also showed increased expression of non-canonical WNT signals (WNT5A and WNT11), which have previously been shown to coordinate emergence of the primitive streak in model animals.^41–43^ To impede non-canonical WNT signalling, we selected the soluble protein secreted frizzled related protein 1 (sFRP1) which will bind all extracellular WNT signals, thereby impeding both canonical and non-canonical pathways. Cells were seeded on patterned 10 kPa substrates of 500 µm diameter with and without soluble SFRP1 (5 µg/ml) for 48h. Immunofluorescence staining of cultures treated with SFRP1 showed maintenance of pluripotency through SOX2, comparative maintenance of E-CAD and no T/BRACHYURY or SOX17 expression (Figure 4B,C,D). In contrast to using the canonical WNT disruptor IWP2, the use of SFRP1 to deplete WNT molecules in the media abolished mes-endodermal differentiation. This suggests the biophysical microenvironment promotes *in vitro* gastrulation through mechanotransduction initiated non-canonical WNT signalling (Figure 4E).

**Figure 4.**
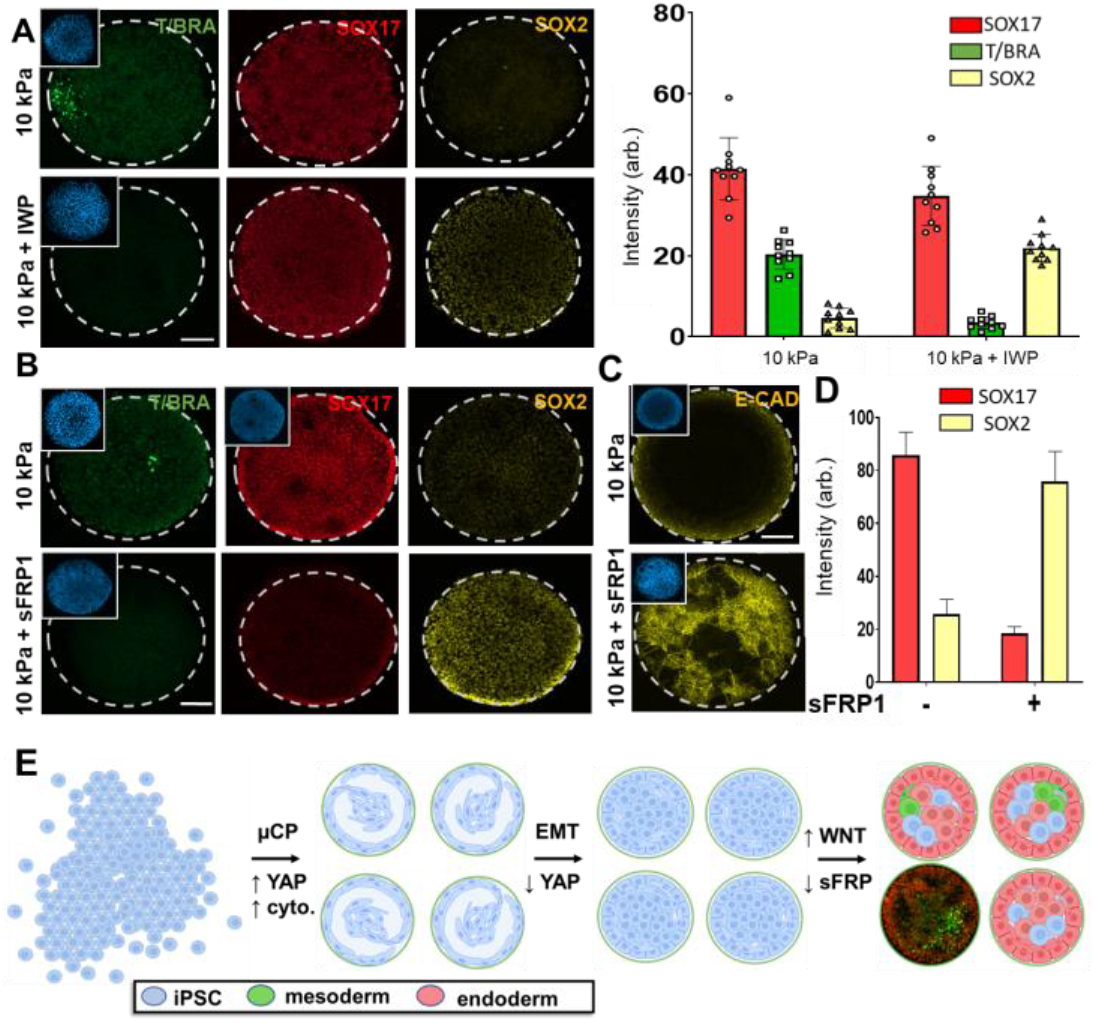
Pharmacological inhibition of canonical and non-canonical Wnt signalling disrupts differentiation. **A)** Immunofluorescence images of micropatterned iPSCs on 10kPa hydrogels with and without canonical WNT inhibition (IWP2). Right – corresponding quantitation. **B)** Immunofluorescence images of micropatterned iPSCs on 10kPa hydrogels with and without total WNT inhibition (sFRP1). **C)** E-cadherin expression with and without sFRP1 treatment. **D)** Quantification of SOX17 and SOX2 expression with and without sFRP1 supplemented media. **E)** Schematic depicting the mechanism where confinement and stiffness promote cytoskeletal tension and YAP activity, EMT and differentation. Scale bar - 100µM

We also probed the effect of small molecule modulators during BMP4 induction by investigating mes-endodermal differentiation on glass and 10kPa patterns in the presence of several modulators of these pathways including the GSKβ antagonist CHIR99021 (CHIR), ALK4/5/7 inhibitor SB431542 (SB) and IWP2 (Figure S10). Treatment with CHIR abrogated spatial patterning in the confined cultures, led to loss of SOX17 expression with a decrease in T/BRACHYURY. Addition of SB along with CHIR results in a further decrease in T/BRACHYURY^+^ cells, demonstrating how combined Nodal and WNT signalling is important for mesodermal differentiation^44^. Addition of SB, either alone or in combination with a WNT activator or inhibitor was expected to abolish endodermal differentiation, as endoderm differentiation is shown to require both WNT and Nodal activity^45^. However, a small population of T/BRACHYURY^+^ cells persisted in CHIR+SB and SB conditions, consistent with WNT signalling being indispensable for mesoderm specification^45^. Blocking both WNT and Nodal in SB+IWP conditions abolished all mes-endodermal differentiation following BMP4 induction, demonstrating the shared role in BMP4 induced gastruloid formation. In the microconfined colonies on 10kPa hydrogels, we observed partial maintenance of SOX17^+^ populations with CHIR and CHIR+SB treatments compared to the sharp decline for colonies on glass in the same conditions (Figure S10).

### Primed colonies form gastruloid-like structures in 3D engineered niches

Since microconfined hiPSCs on gels promote mes-endodermal differentiation without exogenous stimulation, we sought to discern subsequent spatial morphology and organization for gastruloid-like structures harvested from the substrates. After culture for 48 hours, primed embryoids were lifted off the PA surface and individually cultured for 14 days in mTeSR media. To control for the primed conditions, cells were seeded in non-adherent plates to form embryoid bodies (Figure 5A). The embryoid bodies show uniform spherical growth while the gastruloids show an unrestricted structural growth from the periphery over two weeks (Figure 5B). At day 7, we observe a distinct loss of OCT4 expression in the growing gastruloid compared to embryoid, which is further downregulated by day 14 (Figure 5C). Immunostaining the gastruloid for SOX2 (Ectoderm/ Epiblast cells), SOX17 (Endoderm) and T/BRACHYURY (Mesoderm/Primitive Streak) shows loss of OCT4^+^ regions at the interface, with partial SOX2^+^ staining and hotspots of SOX17^+^ and T/BRACHYURY^+^, which is not observed in the embryoids (Figure 5D). The appearance of positional primitive streak-like populations for gastruloids is reminiscent of embryonic gastrulation; however, the magnitude and directionality of the outgrowths vary across samples.

**Figure 5:**
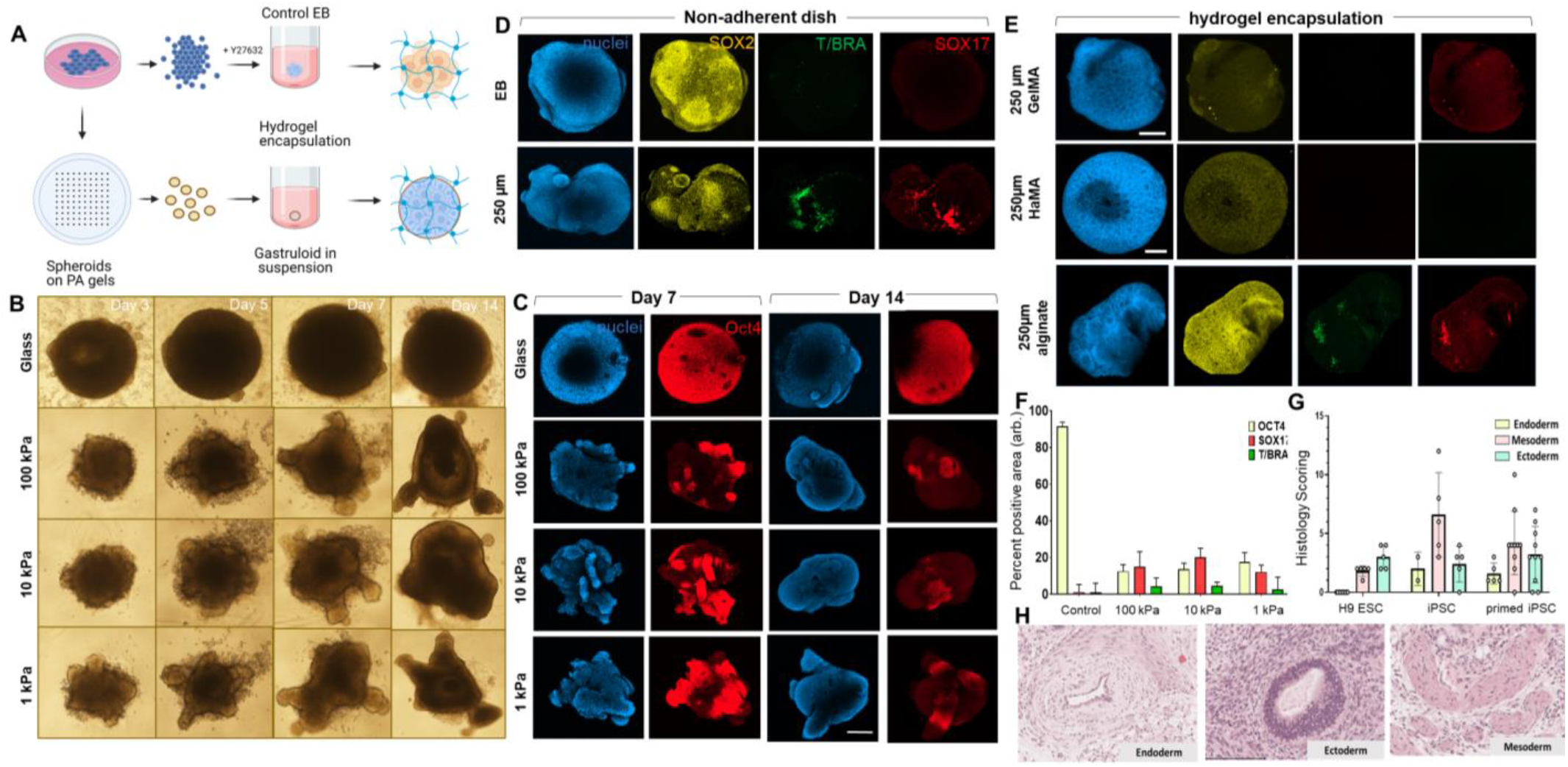
Release from patterns facilitates *in vitro* gastruloids. **A)** Scheme for gastruloid formation and encapsulation within hydrogels. **B)** Brightfield images of embryoid bodies and hydrogel primed gastruloids grown in non-adherent plates. **C)** Immunofluorescence images of embryoid bodies (glass) and gastruloids immunostained for Oct4 at 1 and 2 week culture in non-adherent plates **D)** Immunofluorescence images of embryoid bodies and gastruloids on non-adherent plates and **E)** after encapsulation for 14 days in hydrogel biomaterials. **F)** Percentage of positive area for the OCT4, SOX17 and T/BRACHYURY in embryoid bodies and hydrogel cultured gastruloids. **G)** Histology scoring for H&E scans of teratoma sections from H9 hESCs, hiPSCs and primed hiPSCs. **H)** Representative H&E images for germ layer derivatives identified in the teratoma from (tissue from primed iPSCs shown here). Scale bar - 100µM

Considering the dynamically changing microenvironment surrounding embryonic tissue during gastrulation, we hypothesized that encapsulation of the gastruloids in designer biomaterials that vary ligand presentation, viscoelastic properties, porosity, and diffusion, could be used to guide spatiotemporal patterning. The following hydrogel biomaterials were selected to encapsulate the harvested gastruloid: gelatin methacrylate (GelMA), methacrylated hyaluronic acid (MeHA) and alginate. At day 7, the multi-lobed gastruloids spread out and became more spherical like the embryoid bodies when confined within photocrosslinked gelatin methacrylate (GelMa) or methacrylated hyaluronan (MeHa) with loss of primitive streak hotspots (Figure 5E,F). However, encapsulation within non-covalently stabilised alginate demonstrates elongation of the structure with converging SOX17^+^ and T/BRACHYURY^+^ region, reminiscent of the appearance of a streak like population with mesodermal and endoderm progenitor cells for the initiation of human gastrulation. These results highlight how covalent hydrogel networks (e.g., GelMA, MeHa) may inhibit differentiation and limit further development, while viscoelastic materials that accommodate cell migration and reorganization (e.g., alginate) may be well suited to further guide morphogenesis. Detailed live cell imaging during culture will be necessary to accurately assess growth and morphogenesis within 3D hydrogel matrices. Nevertheless, our results demonstrate how materials can trigger and maintain gastrulation-like states in pluripotent stem cell populations.

To support our *in vitro* differentiation results, we performed a subcutaneous teratoma assay using immunocompromised SCID mice with control pluripotent stem cells (hESC and iPSC), and cells from 10 kPa hydrogels and followed the teratoma growth for 7 weeks. There was high variability in growth across all conditions with no discernible trend in size across starting cell types (Figure S10A). To determin e if the implanted cells undergo differentiation *in vivo*, we performed histology scoring to classify structures associated with different germ layers. We observed duct-like, cartilaginous, bone, loose mesenchyme, smooth muscle, neural rosettes/neuroectoderm, pigmented epithelium, and squamous epithelial-like structures. While the overall histology scoring indicates no significant difference across the cell types (Figure 5G,H), there were subtle variations in the occurrence of specific structures that may be related to the primed mes-endodermal population (Figure S11). Future work will be invested into tracking the differentiated progeny after implantation. However, this is outside of the scope of the present study.

## Discussion

Bioengineered *in vitro* models have become powerful tools to study relationships between biophysical microenvironments and intracellular signalling pathways that align to specify cell fate in populations of pluripotent stem cells^8, 23^. However, these models rely on exogenous addition of soluble factors to trigger differentiation. We have demonstrated how careful selection of cell culture materials can serve to guide morphogenesis and foster conditions to drive *in vitro* gastrulation independent of exogenous chemical induction.

After initial attachment to the micropatterned islands on hydrogels, cells navigated to the regions of elevated stress at the periphery which led to increased cytoskeletal tension and high nuclear YAP localisation. Over time we see enhanced proliferation on the soft hydrogels, which is consistent with previous reports of cells treated with ROCK inhibitor Y27632.^31^ This is in sharp contrast to the populations confined on glass which show uniform adhesion and proliferation. These early aggregates on hydrogels create an epithelial-to-mesenchymal interface, with subsequent differentiation to a mes-endodermal SOX17^+^ T/BRACHYURY^+^ population in the central region. This behaviour of contractile cells at the boundary coordinating EMT has been observed *in vivo*^43^. We propose that hydrogel microconfinement provides analogous initial conditions where the interfacial stress leads to EMT, elevated WNT signalling, and formation of primitive streak-like populations.

The cells lining the perimeter of the colonies co-expressed OCT4, SOX17 and E-CAD, whereas the central part of the colonies showed loss of both OCT4 and E-CAD but with an increase in expression of mes-endodermal markers SOX17 and T/BRACHYURY. The co-expression of OCT4 and SOX17 at the edge suggests partial differentiation consistent with early phases of endoderm specification.^46,47,48^ SOX17 is one of the first transcription factors to be expressed in the inner cell mass^50, 51^ and regulates the dynamics of pluripotency and differentiation.^52^ Moreover, T/BRACHYURY marks the first mesodermal cells that ingress in the primitive streak of a gastrulating embryo.^53, 54^ The patterned phenotypes observed under confinement on hydrogels bears a resemblance to the self-initiation of primitive streak at the onset of gastrulation.

There is considerable evidence to support a central role for biophysical cues directing embryogenesis.^6, 12^ Embryonic epiblast is essentially a layer of epithelial cells tightly connected and packed at the interface as the embryo implants itself on the maternal uterine lining^55^. During gastrulation, the cells at the posterior side of the embryo undergo epithelial-to-mesenchymal transition (EMT)^2, 56^, a signalling gradient arises between BMP, WNT and Nodal pathways, and the primitive streak appears at the same EMT-prone region^1, 57^. *In vivo*, the initiation of EMT precedes the appearance of the first multipotent mes-endodermal progenitors at the primitive streak^56^ which triggers the epiblast layer towards a dynamic, mesenchymal identity at the onset of gastrulation^42^. EMT leads to loss of apical-basal polarity with concurrent nuclear shape changes that contribute to distinct cell-ECM adhesion patterns and motility^58^. These pre-gastrulation events have been shown to be controlled by non-canonical WNT signalling via tight coordinated movements and intercalation of cells to initiate a region prone to EMT^43^ with subsequent mes-endodermal differentiation.

After treatment of microconfined colonies on hydrogels with BMP4, we see significant enhancement of mes-endodermal differentiation with the appearance of multilayered T/BRACHYURY^+^ nodes, the number and position of which is linked to confined area. Transcript analysis indicates that upregulation of mes-endodermal genes is driven by mechanochemical signal transduction that converges with non-canonical WNT signalling. The non-canonical WNT pathways are involved in coordinating EMT^43, 64^ with planar cell polarity pathways guiding the mechanical segregation of cells during gastrulation.^41, 65^ Of the canonical WNT proteins, WNT5A in particular has been linked to regulation of EMT^43^, endodermal differentiation^67^ and regulation of planar cell polarity signalling ^68^, where its loss is responsible for defects in embryological axis elongation and tissue-scale polarity during development.^69, 70^

Mechanotransduction propagates through integrin-mediated adhesion, actomyosin contractility and mitogen activated protein kinase signalling, where numerous downstream effectors are employed by both matrix engagement and soluble morphogens^22^. Here we showed how adhesion to deformable hydrogels leads to enhanced cytoskeletal tension and YAP activity at the boundary which triggers EMT across the colony. Once EMT is underway, differentiation is coordinated through complementary paracrine/autocrine signalling, like the planar cell polarity pathway involving both WNT5A and WNT11. WNT signalling controls differentiation during vertebrate development, and the planar cell polarity serves as a route for organising tissue patterning^24, 25, 42, 71^. After treating our cultures with small molecule inhibitors for WNT/β-catenin and TGFβ pathways we see some moderate attenuation of mes-endodermal differentiation. The small molecule inhibitor IWP targets canonical WNT signalling involving β-catenin translocation, and does not impede non-canonical pathways. IWP treatment inhibits the expression of mesoderm marker T/BRACHYURY but leads to no change in the expression of endoderm marker SOX17. In contrast, treatment of our cultures with SFRP1, which will inhibit all WNT signalling in the extracellular space, leads to complete abolishment of both markers associated with a primitive streak-like populations. Since differentiation still occurs with inhibition of canonical WNT signalling, this result demonstrates a central role for non-canonical WNTs in coordinating *in vitro* gastrulation.

Recently, there have been several reports of pluripotent stem cell colonies patterned on glass substrates for *in vitro* models of neuroectoderm^62^, cardiac tissue^18^ and coordination of pre-streak patterning.^63^ Across all of these studies, geometric confinement was shown to guide cellular assembly, where triggering differentiation with soluble morphogens would lead to biomimetic forms with distinct functions. Weaver and colleagues showed how soft substrates will bias pluripotent stem cells towards mesodermal differentiation following BMP4 induction, providing evidence for the importance of matrix mechanics during lineage determination^15^. Previously, we showed how combining deformable matrices with geometric confinement promotes EMT via mechanotransduction in populations of cancer cells leading to epigenetic reprogramming and emergence of a stem cell phenotype.^59,60,61^ Here we show that confining pluripotent stem cells on deformable hydrogels serves to guide cellular assembly and partitioning, thereby providing the right context to trigger EMT and catalyse morphogenetic events that mirror *in vivo* processes of differentiation.

The ability to foster gastrulation-like events through the properties of the culture substrate alone opens opportunities to study the relationships between the biophysical microenvironment and early embryogenesis. Colonies that have been primed for two days in microconfinement were released from the substrate and cultured in suspension to see if further patterning would occur in standard conditions for embryoid body growth. Compared to control embryoid bodies which exhibit a uniform morphology with robust staining of pluripotency markers, the primed colonies exhibited clear regions of mes-endodermal identity which further evolved into lobed structures reminiscent of embryo patterning over the course of 14 days. The morphology of these gastruloids show similarities to approaches involving small molecule stimulation of embryoid bodies^72^. Encapsulation of the gastruloids within a library of hydrogel biomaterials demonstrates materials parameters that promote and prevent continued lineage specification. These finding underscore the importance of materials in nurturing the differentiated phenotype during *in vitro* gastrulation, which draws parallels to the role of the extracellular matrix in orchestrating embryogenesis *in vivo*.

## Conclusions

Modelling human gastrulation *in vitro* through materials properties without exogenous biochemical induction raises numerous opportunities for probing fundamental stages in embryogenesis. The precise control of gastrulation *in vivo* is afforded by tight coordination of soluble signals with feedback from the biophysical microenvironment. Using microengineered hydrogel substrates, we demonstrate a method where controlling the biophysical microenvironment poises a population of pluripotent stem cells to undergo morphogenesis, with integration of soluble signals to orchestrate gastrulation-like processes. These findings provide a new avenue for probing the biophysical and biochemical basis of embryogenesis, and a tool to model development for fundamental biology and translational endeavours.

## Methods

### Cell culture and Maintenance

Hepatic fibroblast derived iPS line ATCC-HYS0103 Human Induced Pluripotent Stem (IPS) Cells were purchased directly from the vendor. hESC-Qualified Matrigel (Corning 354277) Matrigel was used coat culture dishes for feeder-free expansion of induced pluripotent stem cells and were routinely cultured and maintained in mTeSR™1 (STEMCELL Technologies, 85850) in a humidified incubator at 37 °C with 5% CO_2_. Cells were passaged once a week using selective dissociation reagent ReLeSR™ (STEMCELL Technologies 05872) and seeded using the cell aggregate counting method described in ‘Plating Human ES and iPS Cells Using the Cell Aggregate Count Method’ (Appendix 1) from the STEMCELL Technologies-‘Maintenance of Human Pluripotent Stem Cells in mTeSR™1’ technical handbook to assess the size of aggregates and seed them in low, medium or high densities, as described. All cryopreservation of cells was performed in CryoStor® CS10 (STEMCELL Technologies 07930) freezing media.

For the glass controls in the experiments – sterile rh-Vitronectin (Gibco, Life Technologies, A14700) was used to coat glass cover slips. In case of all experiments, cells were dissociated into a single cell suspension using StemPro™ Accutase™ (Gibco A1110501) cell dissociation reagent to facilitate cell counting. In case of single cell dissociation, cells were always seeded with 10uM Rock inhibitor Y27632 (ATCC® ACS-3030™).

### Preparation of PA coated cover slips

Hydrogel based substrates were prepared using chemically modified polyacrylamide and soft lithography. Round glass cover slips (18mm Diameter) were sonicated and individually placed in a 12-well tissue culture polystyrene plate, treated with 0.5% 3-Aminopropyl triethoxysilane (APTS) (Sigma Aldrich A3648) for 3 minutes then with 0.5% Glutaraldehyde (Sigma Alrich G6257) for 30 minutes. The cover slips are thoroughly air dried with the treated surface up. For the polyacrylamide hydrogel coating, 40% solution of Acrylamide (Sigma Alrich A3553) and 2% solution of Bisacrylamide (Sigma Aldrich 146072) were prepared in distilled water. Solutions pertaining to various elastic modulus were prepared as described in the table ^73^. The stiffness solution was sandwiched between a hydrophobic glass slide and the treated cover slip. 10 % Ammonium Persulfate (APS) and Tetramethylethylenediamine (TEMED) (Sigma Aldrich, 1.10732) were used for polymerization in a covered, moist environment. Cover slips were carefully picked up then treated with Hydrazine hydrate 100% (Acros organics 196715000) for up to 1 hour to convert amide groups in polyacrylamide to reactive hydrazide groups and then 1 hour incubation in a 5% solution of Glacial acetic acid is done before patterning. DI washes performed between each chemical treatment.

### Microcontact Patterning of PA coated cover slips

For microcontact patterning, polydimethylsiloxane (PDMS, Polysciences, Inc.) stamps for 250u and 500u diameter circles were prepared by polymerization upon a patterned master of photoresist (SU-8, MicroChem) created using UV photolithography using a laser printed mask. Sodium periodate (Univar 695-100G) solution was used to yield free aldehydes in the proteins used for micropatterning. For assisting adhesion of iPS cells, recombinant human Vitronectin (Gibco A14700) was used at a final concentration of 25ug/ml. The solution is applied to the patterned or non-patterned PDMS stamps for 30 minutes, air dried and then applied to the air-dried PA surface. The stamps are removed after 15 seconds and the patterned cover slips are sterilized by transferring to a sterile 12-well TCP dish inside BSCII and 3x Sterile DPBS wash, followed by 12 minutes of UV exposure and then stored in 4^◦^C soaked in a 2% Penicillin/Streptomycin + DPBS solution for minimum 6 hours. The soaking solution to be removed and warm expansion media to be added to the cover slips before seeding the cells.

### Microcontact patterning of glass cover slips

The same PDMS stamps were used to assisnt with physical adsorption of protein on clean, dry glass cover slips. The glass cover slips were sonicated in 100% ethanol and cleaned in a plasma cleaner (PlasmaFlo, Harrick Plasma) to remove all residue. Vitronectin was dissolved in PBS at the final concentration of 25µg/ml and applied on clean stamps 30mins/37^◦^C. The stamps were rinsed, dried and applied on clean cover slips for 5-7 minutes allowing for protein adsorption on defined islands. The patterned glass cover slips were sterilised before cell culture and used within 24 hours of preparation.

### Cell seeding on the micropatterned cover slips

The cell culture dishes are monitored for confluency before starting the experiment. All experiments were performed using in mTeSR™1 (STEMCELL Technologies, 85850). The cells are dissociated using warm StemPro™ Accutase™ (Gibco A1110501) cell dissociation reagent for 6-7 minutes. Cells were counted using a haemocytometer and seeded in mTeSR™1 (STEMCELL Technologies, 85850) and a density of 5×10^5^ cells/ml as a 1ml/well solution in a 12-well culture dish with 10uM Rock inhibitor Y27632 (ATCC® ACS-3030™). The medium was replaced with media without Y-27632 at 24h for 10 and 100kPa substrates, and 5uM Y-27632 in case of 1 kPa substrates (This was completely removed at 36h). The cells were allowed to grow on the 4 substrate groups for 48h.

For BMP4 induction experiments, same seeding method was used, fresh media supplemented with rhBMP4 (STEMCELL Technologies, 78211) at a final concentration of 50ng/ml on the 4 substrate groups for 48h.

For small molecule inhibitor treatments-cells were supplemented with the following concentration of each small molecule at 6h of seeding– CHIR99021 (3uM), SB431542 (5uM), IWP-2 (2.5uM), sFRP1 (5ug/ml).

### Immunocytochemistry – fixation and staining

Cell media was replaced with 4% solution of Pierce™ 16% Formaldehyde (w/v), Methanol-free (Thermo Fisher Scientific, 2890) diluted in 1X PBS, for 30 minutes at RT. All washes were performed using Gibco™ DMEM, no phenol red (Gibco, 31053028). This was done because any contact with PBS would readily detach all cells from the patterned PA surfaces. 0.1% solution of Triton X-100 in PBS (v/v) and 3% Bovine Serum Albumin solution (Sigma Aldrich, A3803) in PBS (w/v) was used for permeabilization and blocking. Primary incubation 4^◦^C overnight/secondary incubation for 1 hour at RT in dark. Coverslips were mounted using ProLong™ Gold Antifade Mountant (Thermo Fisher Scientific, P36930) which contains a DAPI stain.

### 3D Spheroid culture and Immunostaining

At 48h, the media on the PA gels was removed, and PBS washes were performed 2-3 times, whilst collecting the PBS after each wash in a separate labelled tube for each condition. As mentioned above, PBS washes would readily dissociate all colonies, but the spheroid structure of the colonies are maintained. 50ul of warm mTeSR™1 was added to a 96-well Low adherence U-bottom dish. Individual spheroids are picked up and deposited in the 96 well. After checking each well for only one spheroid each, the wells are topped up with 100ul of media. The spheroids are grown for 14 days, with media change done every other day. Similarly, spheroids were deposited in the biomaterials and cultured for 7 days. 5000-15000 cells deposited in each well with Y27632 supplemented for controls, cultured 2 days before being deposited in the biomaterials.

For fixation, a cut 200ul tip is used to pick up the spheroid and transferred to a glass-bottom 96 well plate and fixed with 4% PFA (Thermo Fisher Scientific, 2890) for 24 hours. 2x PBS washes of 1 hour each was done followed by 0.2% solution of Triton X-100 for 2 hours and blocking for 1 hour. Primary and secondary antibody+Hoescht staining was performed for 24h each on a slow rocker at RT with 2×1 hour PBS washes in between.

### *In vivo* teratoma analysis

The intact spheroids were collected off the 10kPa PA gels via PBS washes as described above. The spheroids were allowed to settle in the PBS for 10 minutes at RT, supernatant was removed, and the spheroids were resuspended in 100µl Matrigel-GFR (Corning 354230). ATCC hiPSCs and H9 hESCs were used as controls. This was injected in immunocompromised SCID mice subcutaneously and was observed for growth for the next 7 weeks. The teratomas were collected in ice-cold PBS, fixed in 10% neutral-buffered formalin solution for 3 days, and transferred in 70% ethanol for 3 days. Paraffin embedded sectioning, mounting and haematoxylin/eosin staining was performed for all sections. Imaging of slides was done using Leica Aperio XT slide scanner. Histology scoring was plotted using Graphpad Prism.

### RNA isolation and qRT-PCR

For RNA isolation, cells on all four groups of substrates were dissociated using StemPro™ Accutase™ (Gibco A1110501) for 5 minutes. Cell pellets were washed with RT PBS and then with chilled PBS while being centrifuged at 4C. All existing PBS is removed, cell pellets were snap-frozen using liquid nitrogen and stored in −80. RNA isolation is performed using RNeasy Mini Kit (Qiagen 74104) as per kit instructions. cDNA prep was performed using RT2 First Strand Kit (Qiagen 330401) as per kit instructions. The cDNA was added to RT² SYBR Green ROX qPCR Mastermix (Qiagen 330520) and the solution was added to a Custom RT2 PCR Array 384-well plate and qPCR program was run using QuantStudio™ 7 Flex Real-Time PCR System (Applied Biosystems 4485701). Data analysis was performed on https://geneglobe.qiagen.com/us/analyze. Heatmaps prepared on GraphPad Prism and Morpheus.

### Confocal imaging and data quantification

The fluorescence imaging was performed on a Zeiss LSM780 or a Zeiss LSM800 confocal microscope, normally using 10x and 20x objectives and images were acquired using the Zen Pro imaging software from Zeiss. Image analysis was performed using FIJI (Fiji is just ImageJ) software, and extra plugins were downloaded when required. Nuclear characteristics analysis was performed using CellProfiler Cell image analysis software from Broad Institute. Yap localisation and some nuclear measurements were performed using a MATLAB based GUI prepared for our data. Data sets were compiled in Microsoft Excel and graphs preparation and statistical analysis was done using GraphPad Prism.

**Table:1.**
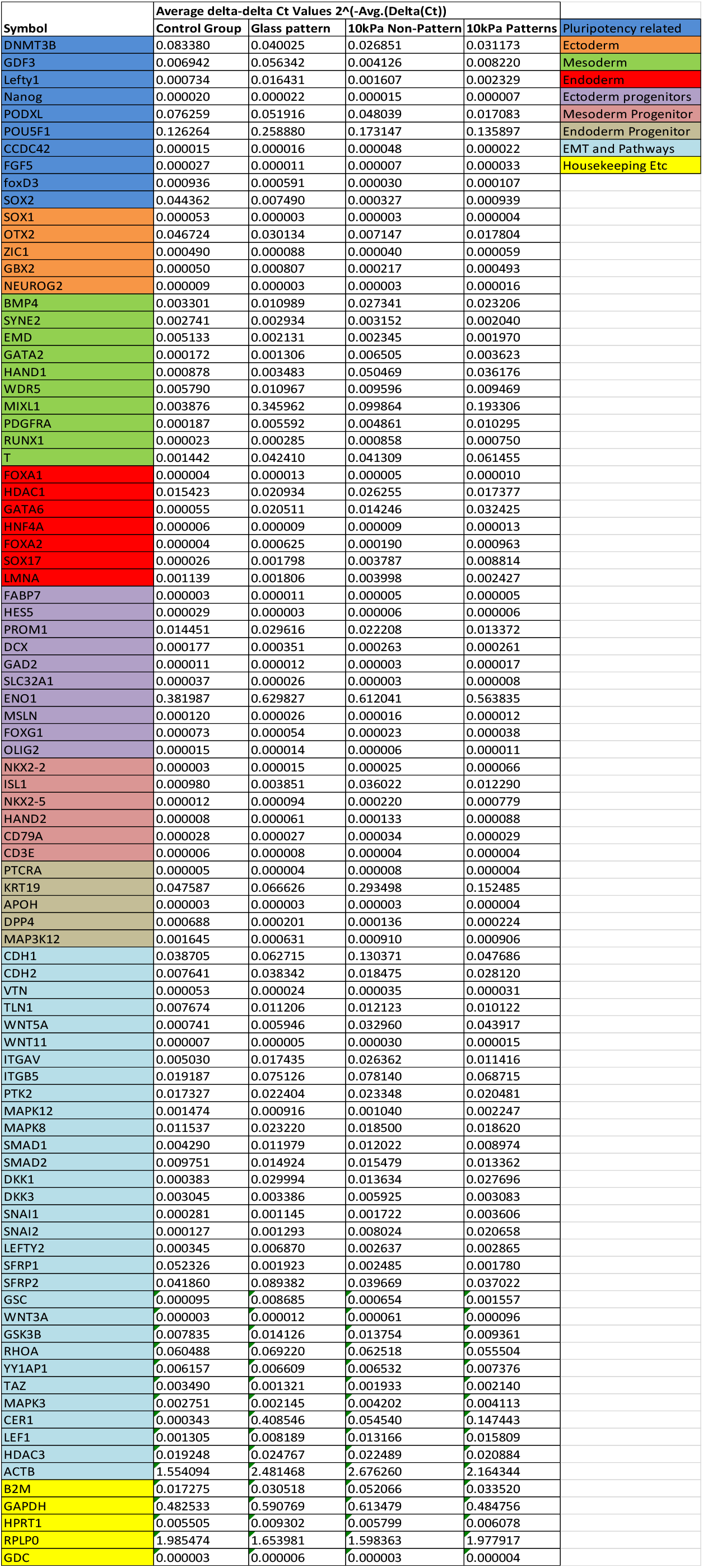
Raw data -Gene list+Av double delta Ct values (2^ ΔCt) for +BMP4 gene expression analysis

**Table 2:**
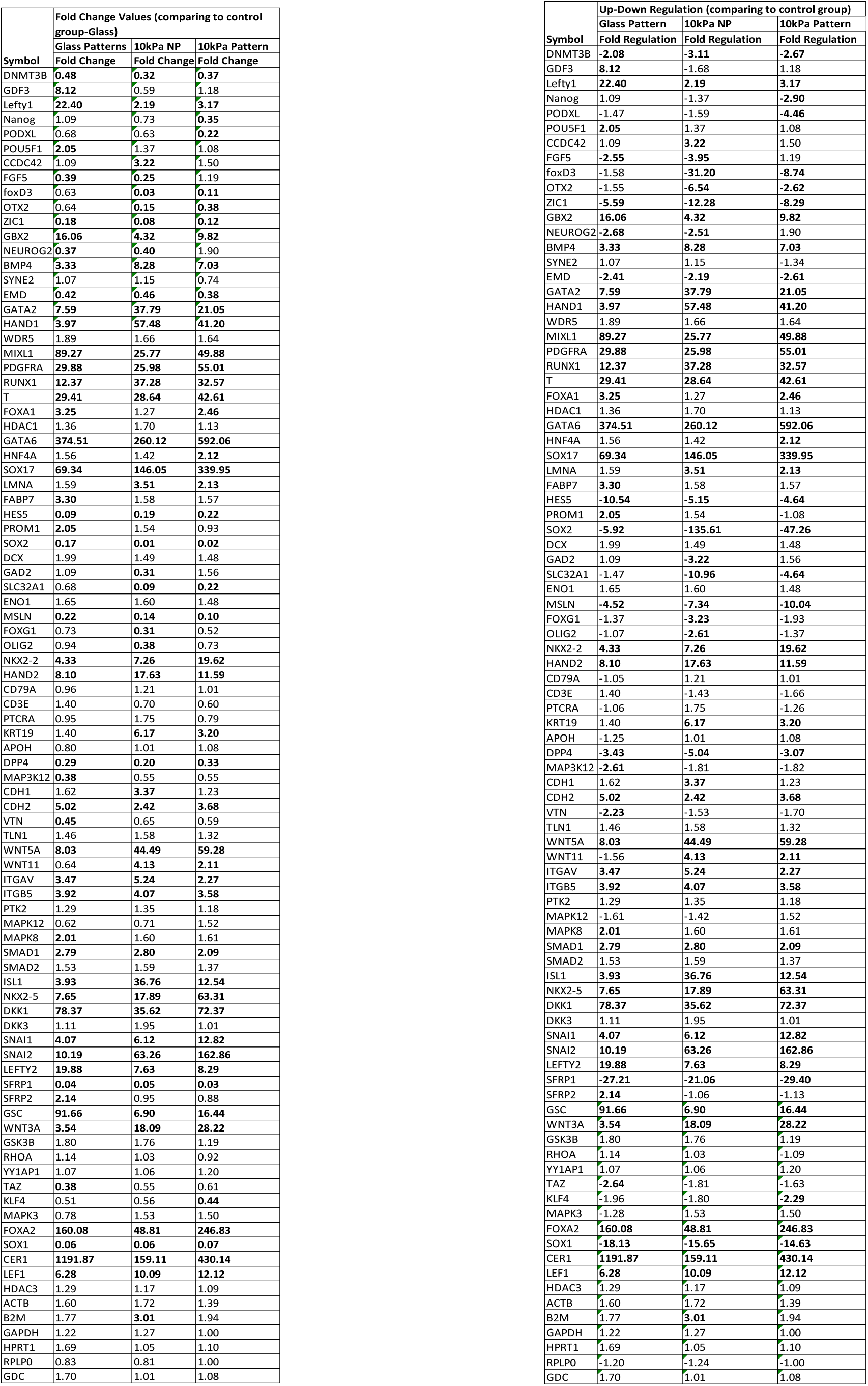
Raw data -Gene list+fold-change and fold regulation values for +BMP4 gene expression analysis

**Table 3:** Antibody Information:

| Antibody name | Cat. No | Produced in | Dilution used |
| --- | --- | --- | --- |
| Oct4 | Goat Polyclonal Anti-Oct4 antibody (ab27985) | Goat | 1:500 |
| Nanog | Nanog Monoclonal Antibody (hNanog.2), eBioscience 14-5768-82 | Mouse | 1:400 |

Differentiation markers:
| Antibody name | Cat. No | Dilution Used |
| --- | --- | --- |
| Sox17 | RDSAF1924 R&D Systems Human SOX17 Affinity Purified Goat Polyclonal Ab | 1:300 |
| T/Brachyury | ab209665 Rabbit monoclonal [EPR18113] to Brachyury | 1:500 |
| Sox2 | ab79351 Mouse monoclonal [9-9-3] to SOX2 | 1:500 |
| E-Cadherin | Mouse monoclonal [M168] to E Cadherin ab76055 | 1:500 |
| FoxA2 | Mouse monoclonal [7E6] to FOXA2 ab60721 | 1:500 |
| Sox1 | SOX1 Recombinant Rabbit Monoclonal Antibody MA5-32447 | 1:500 |
| Snail | SNAIL Rabbit Monoclonal Antibody (F.31.8) MA5-14801 | 1:400 |

2<sup>nd</sup> antibodies
| Antibody name | Cat. No | Dilution Used |
| --- | --- | --- |
| Anti-goat 647 | ab150131 Donkey Anti-Goat IgG H&L (Alexa Fluor® 647) | 1:400-1:600 |
| Anti-rabbit 488 | Donkey anti-Rabbit IgG (H+L) Highly Cross-Adsorbed Secondary Antibody, Alexa Fluor 488 A-21206 | 1:400-1:600 |
| Anti-Mouse 555 | Goat anti-Mouse IgG (H+L) Highly Cross-Adsorbed Secondary Antibody, Alexa Fluor 555 A-21424 | 1:400-1:600 |

## Author Contribution

PS, JP and KK developed the ideas. PS, SR, JI, TM, SN, PJ, AY, EP and VC performed the experiments and analysed the data. All authors contributed to writing the manuscript.

## Acknowledgements

This work was supported through funding from the Australian Research Council Grant FT180100417. The authors acknowledge the help and support of staff at the Biomedical Imaging Facility and the Biological Specimen Preparation Laboratory of the UNSW Mark Wainwright Analytical Centre. We would also like to thank Dr. Pierre Osteill for helpful conversations and guidance over the course of this work. Icons were generated on Biorender.com.

## Supplementary Information

**Figure S1:**
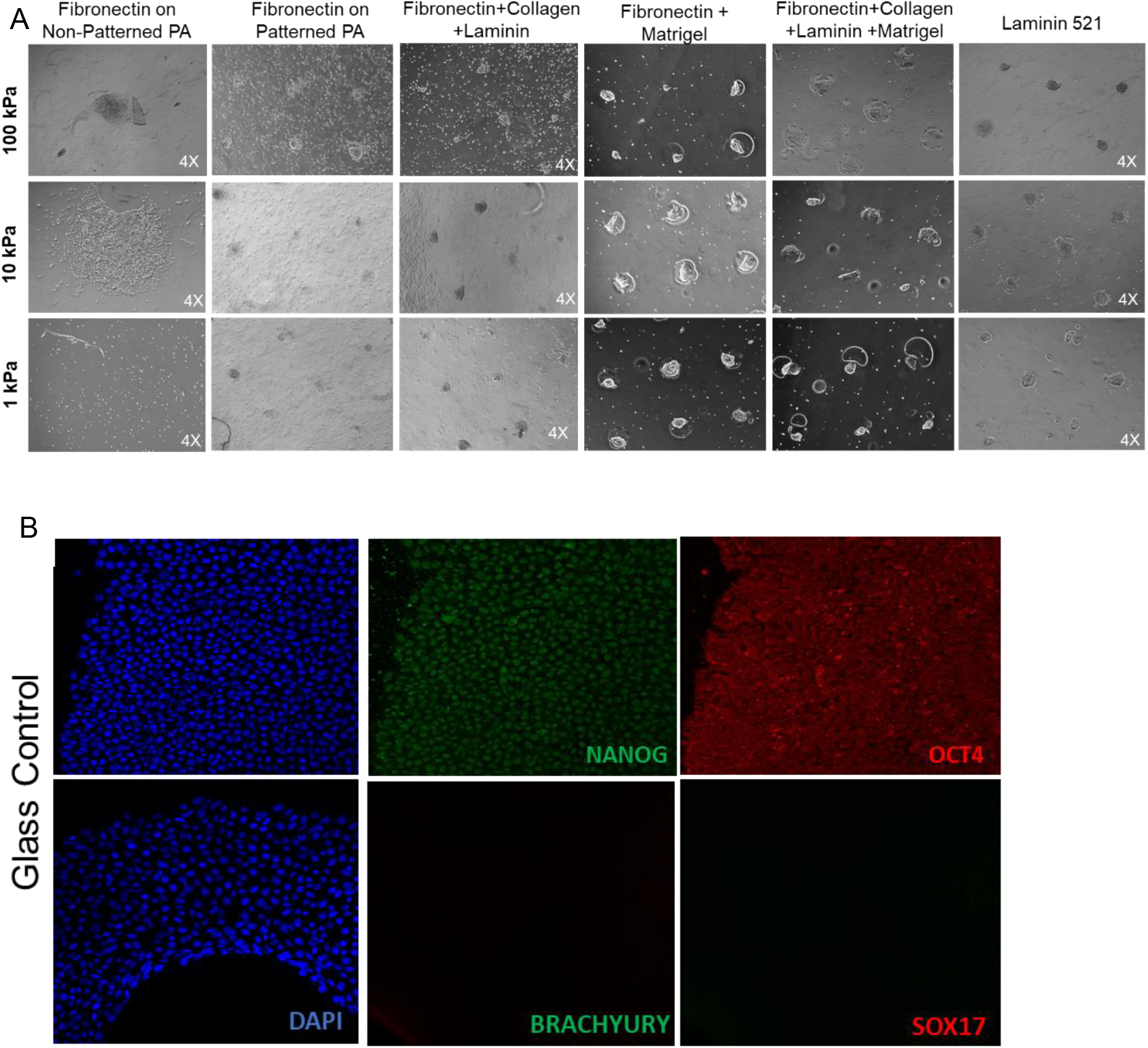
A) The panel of various ECM proteins tested for cell adhesion on polyacrylamide hydrogels, shows inadequate adhesion and proliferation in all combinations at 48 hours. B) ATCC hiPSC characterisation Scale bars: 100µm

**Figure S2:**
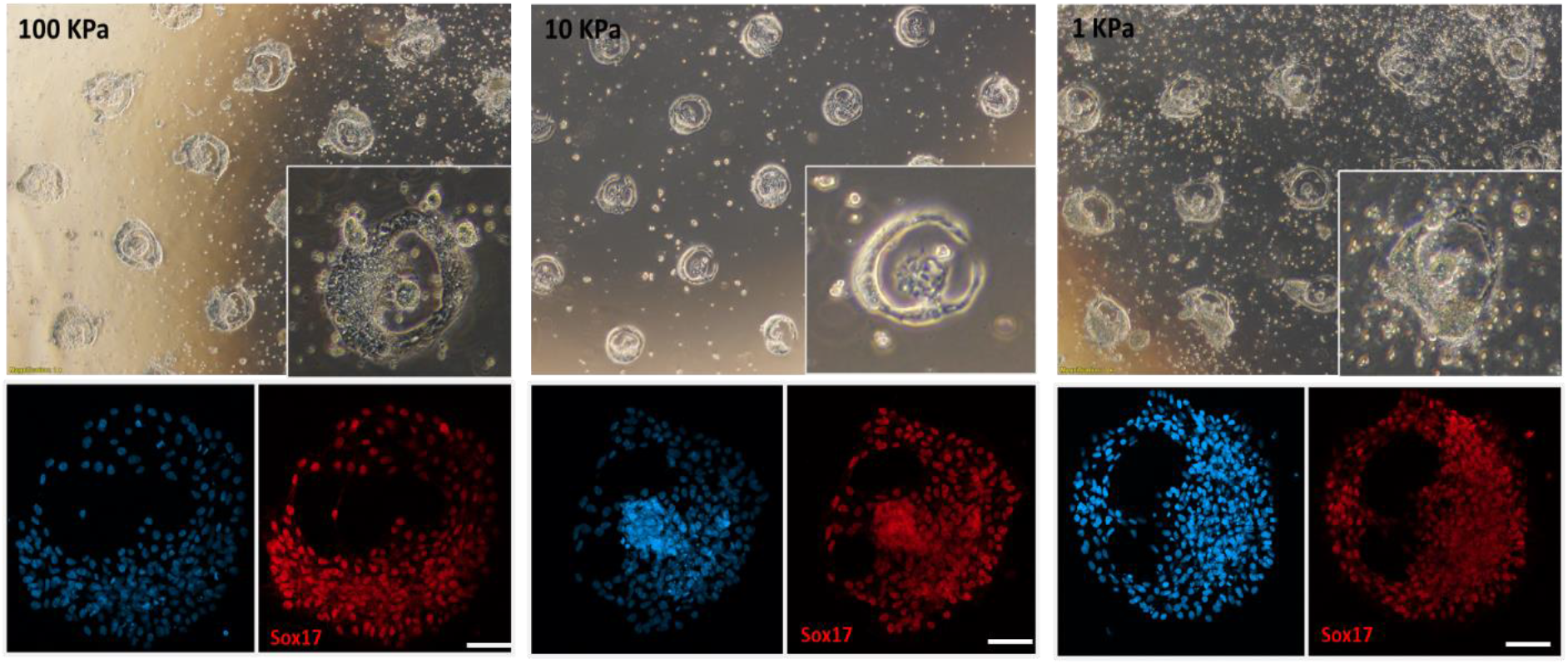
Brightfield and IF images of hiPSCs seeded on PA gel substrates with 250um circular patterns at 12 hours. The cells seem to circle the colony boundary before depositing towards the colony centre. The cells were seen to be of Sox17+ endodermal identity as early as 12h post seeding. Scale bars: 50µm

**Figure S3.**
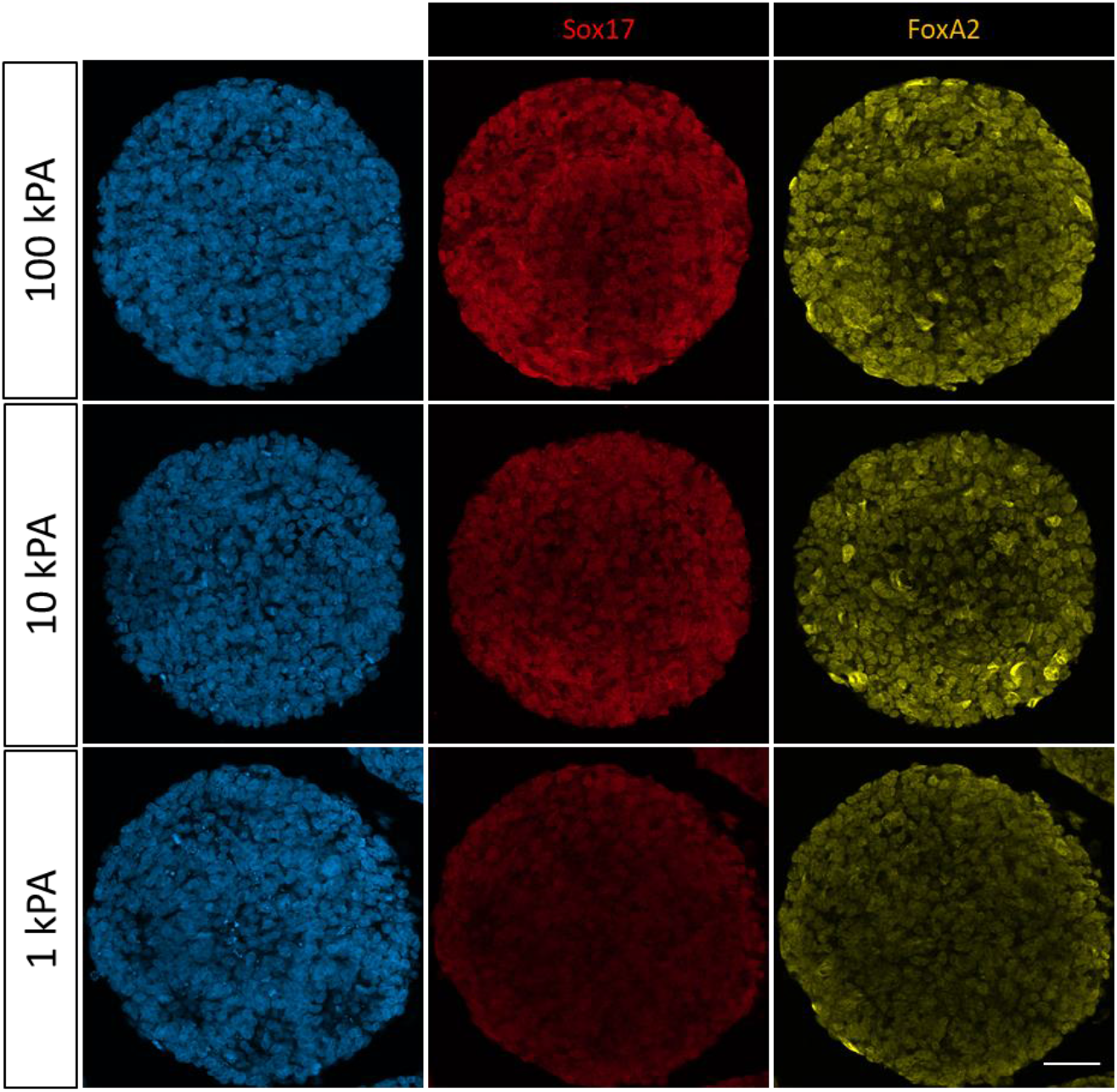
Colonies 250uM circles on PA gels positive for Sox17 as well as FoxA2 Scale bar – 50µm

**Figure S4:**
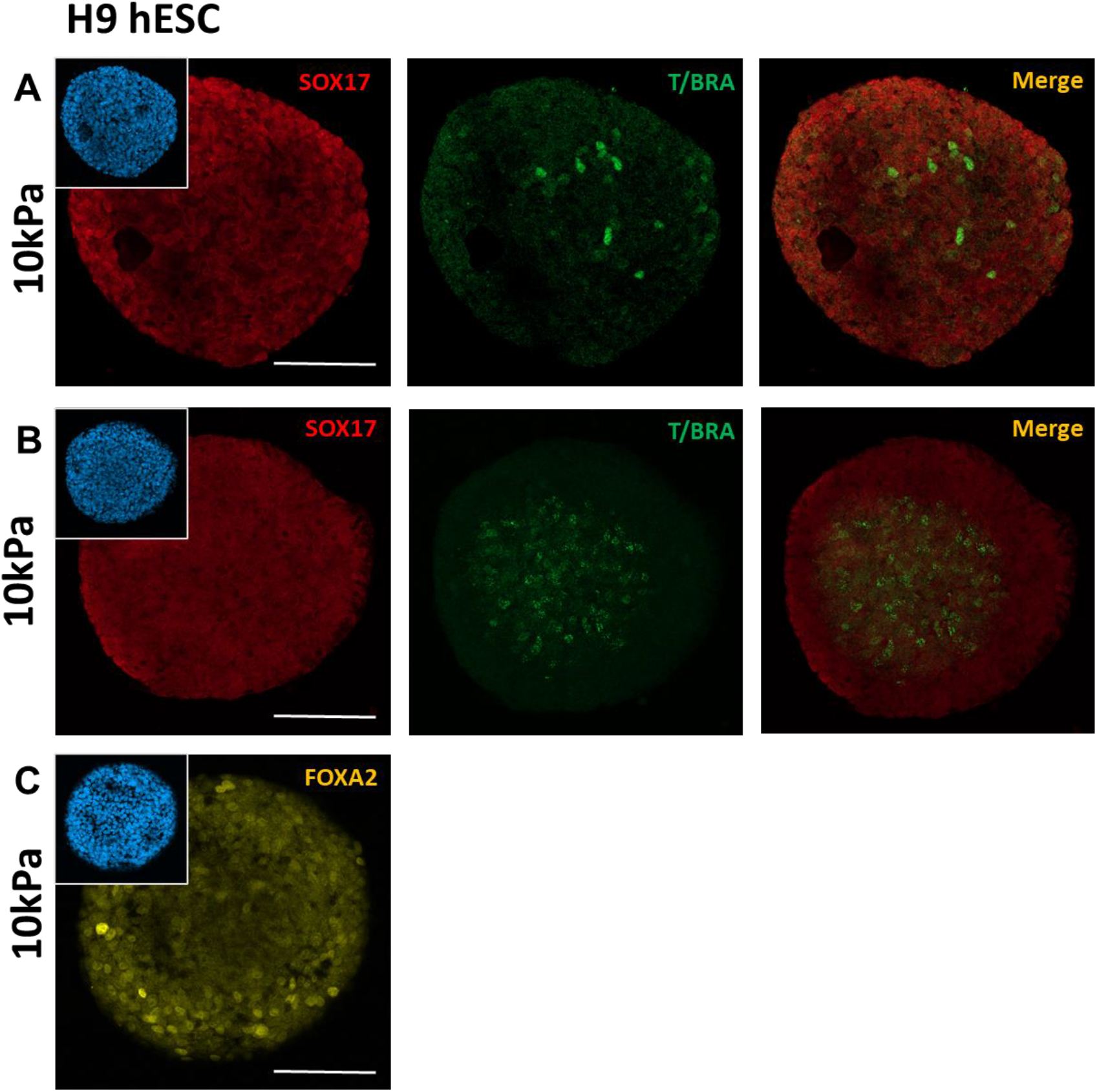
Immunofluorescent images of H9 hESCs seeded on 10kPa PA gel substrates with 250um circular patterns at 48 hours. A) The hESCs assumed an endodermal SOX17+ identity with a mesodermal T/BRACHYURY+ cluster within. B) In absence of a mesodermal cluster, nuclear puncta were observed in colony centres in response to T/BRACHYURY staining. C) Majority of SOX17+ cells co-expressed endodermal marker FOXA2 Scale bars: 100µm

**Figure S5:**
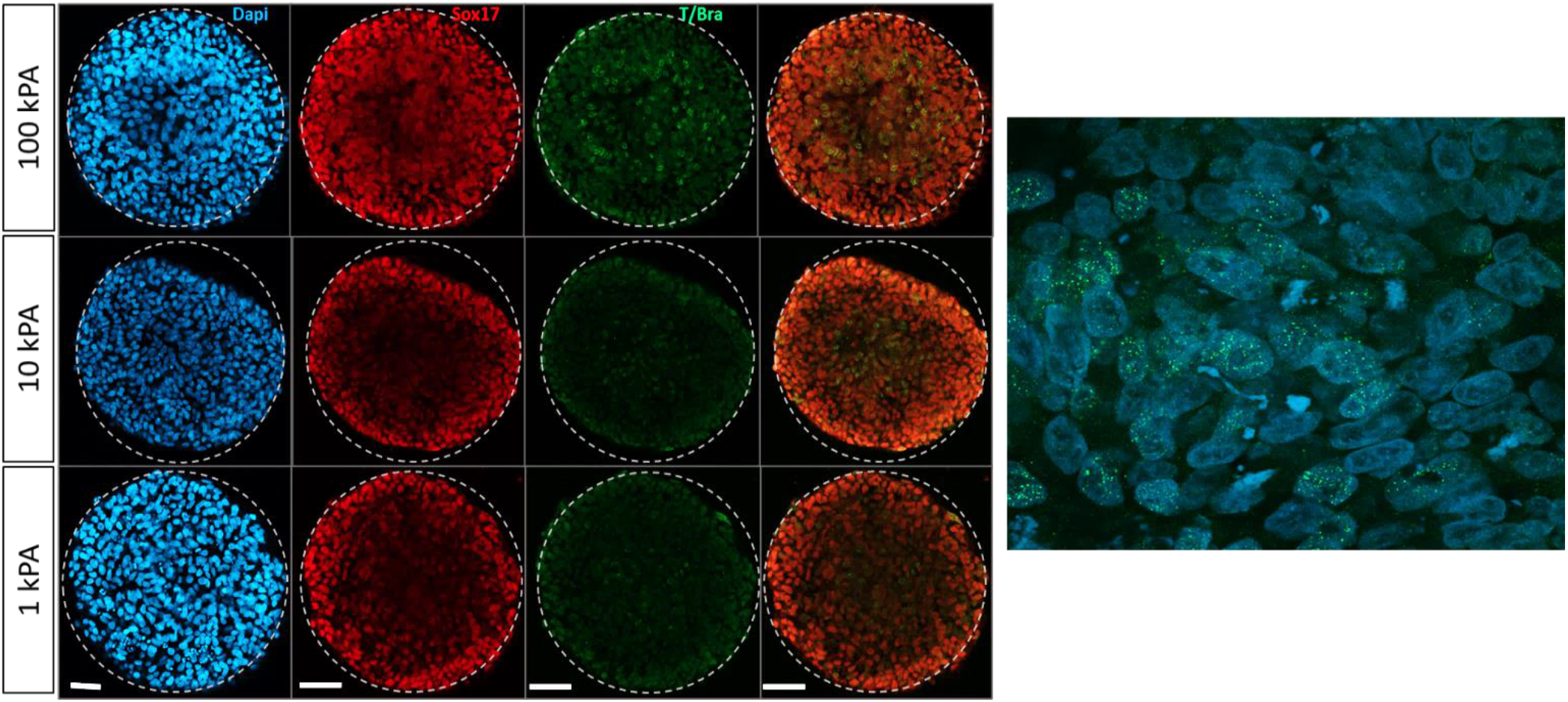
Stiffness and geometrical confinement create distinct nuclear punctates for Mesoderm/Primitive Streak marker T/Brachyury, regularly observed in the cells towards the centre of the hiPSCs colonies seeded on PA hydrogel substrates with circular patterns of 500µm diameter. Right – 63x image to show nuclear puncta. Scale bars – 100µm

**Figure S6:**
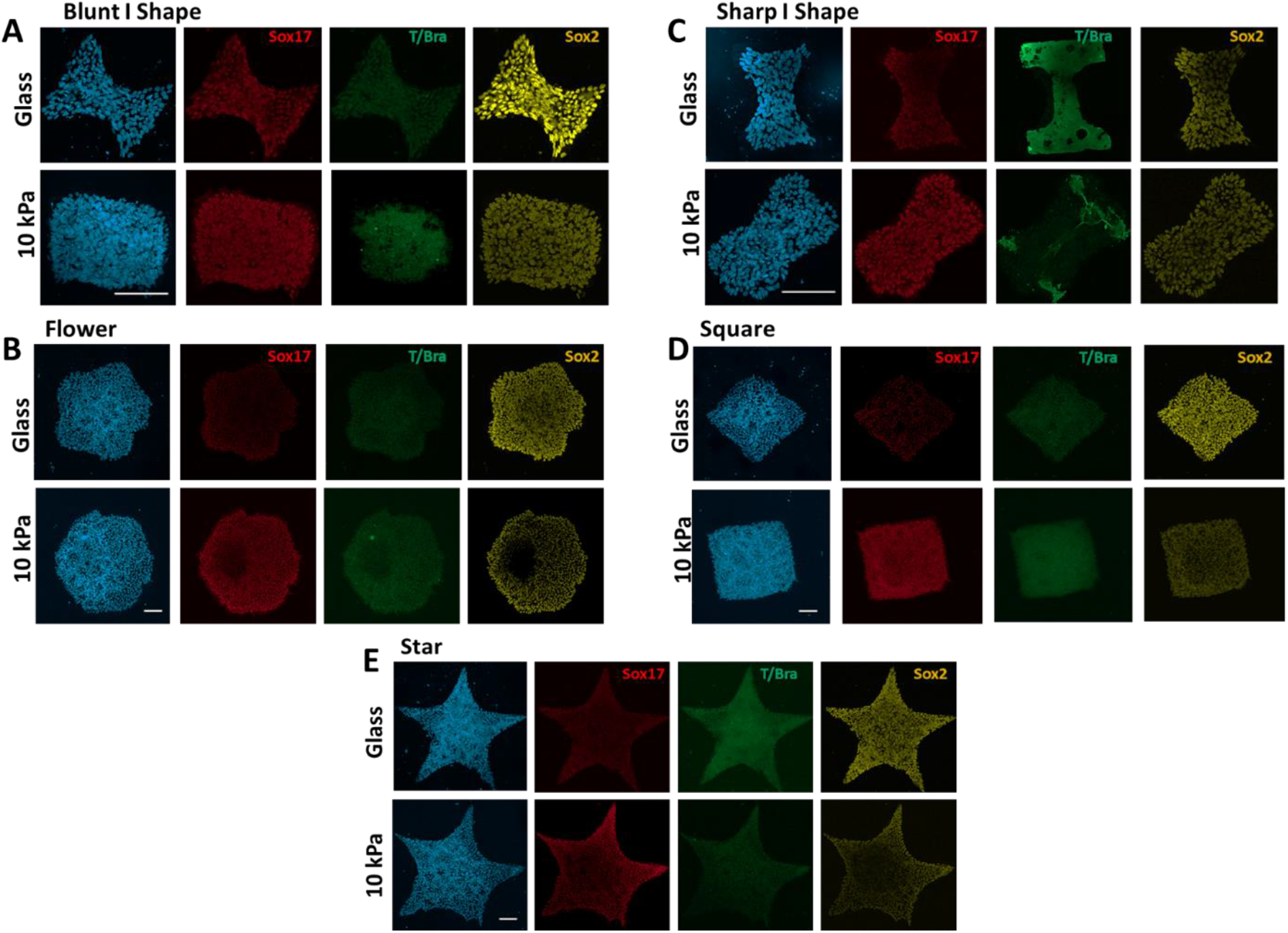
Immunofluorescent images of ATCC hiPSCs seeded on glass 10kPa PA gel substrates patterned using various shapes at 48 hours. A) Blunt ‘I’ shape B) Flower C) Sharp ‘I’ shape D) Square, and E) Star. Scale bars: 100µm

**Figure S7:**
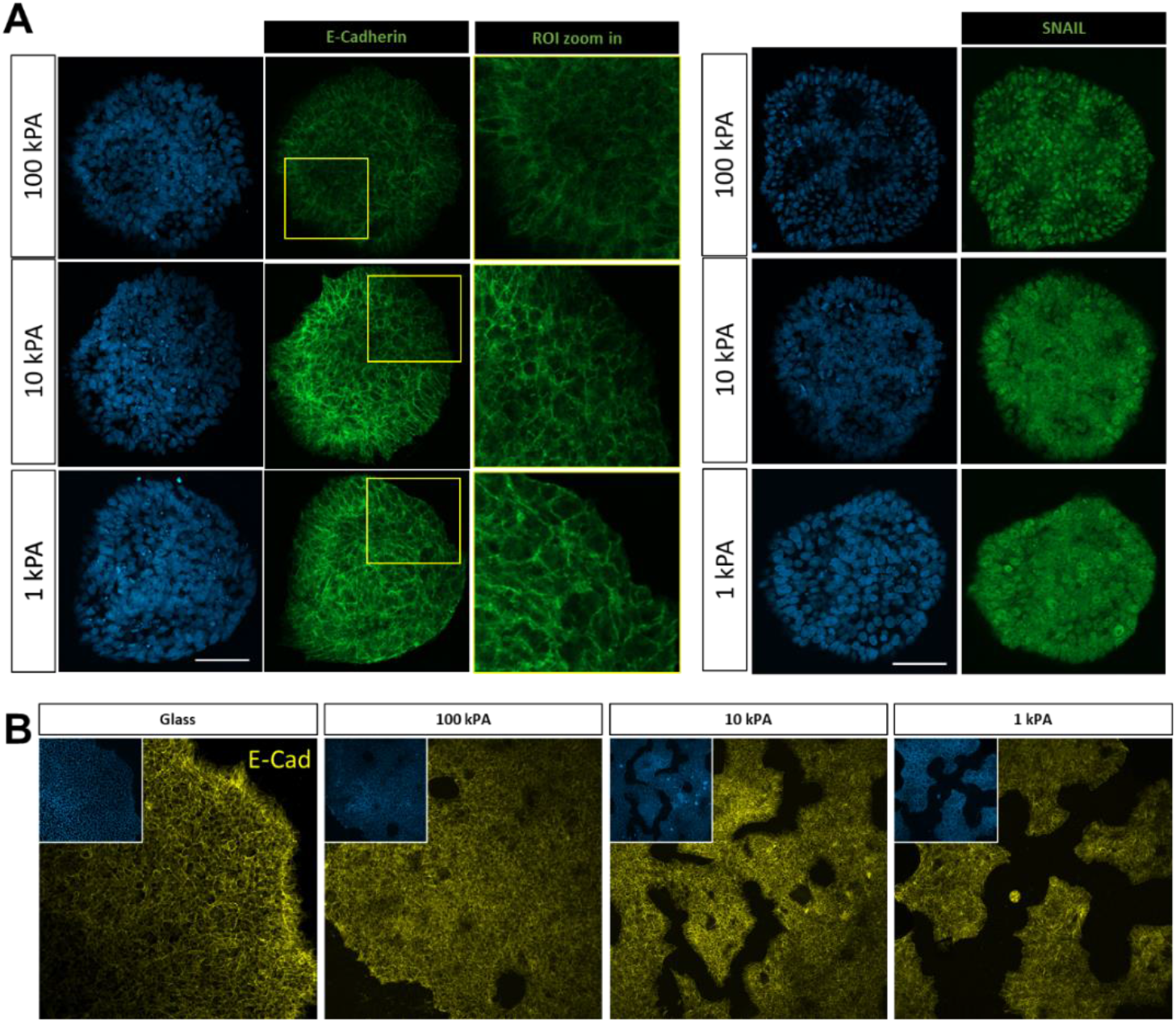
EMT Signifiers on non-BMP4 experiments A) Discontinuous regions of E-Cadherin and Expression of EMT marker snail in the 250um diameter colonies: signifies occurrence of EMT: Significant discontinuity and cytoplasmic localization of E-Cadherin in the colonies on the PA hydrogel substrates. Global Expression of SNAIL on the 3 stiffness substrates, however the clearest expression is observed on 100kPa. Both these act as an important signifier of EMT. B) E-cad expression on glass and Non-Patterned gels Scale bars: 100µm

**Figure S8:**
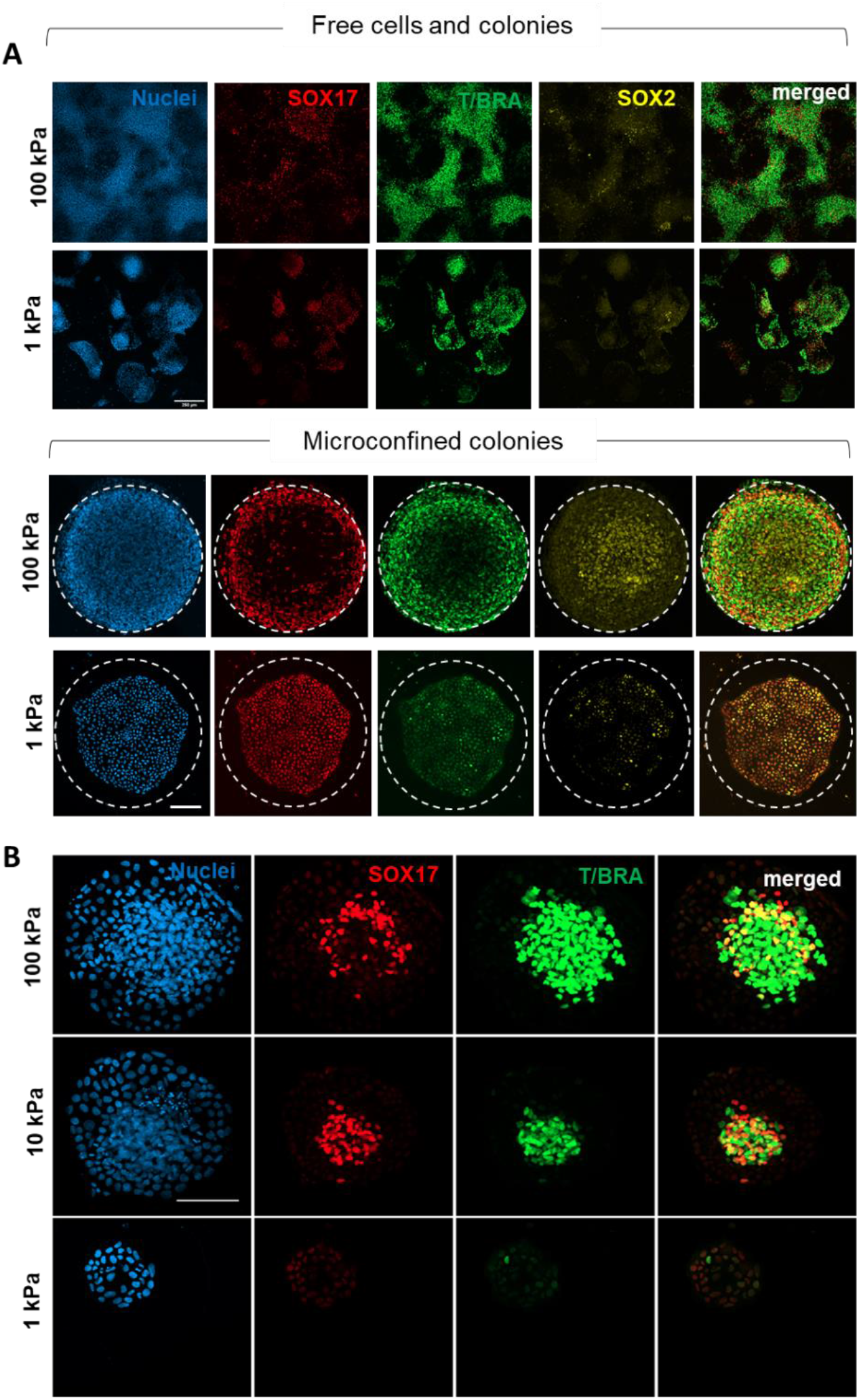
A) Immunofluorescence markers for mes-endodermal differentiation for iPSCs cultured on 1 kPa and 100 kPa hydrogels. B) Immunofluorescence markers for mes-endodermal differentiation for iPSCs cultured on 250 µm^2^ islands. Scale bars: 100µm

**Figure S9:**
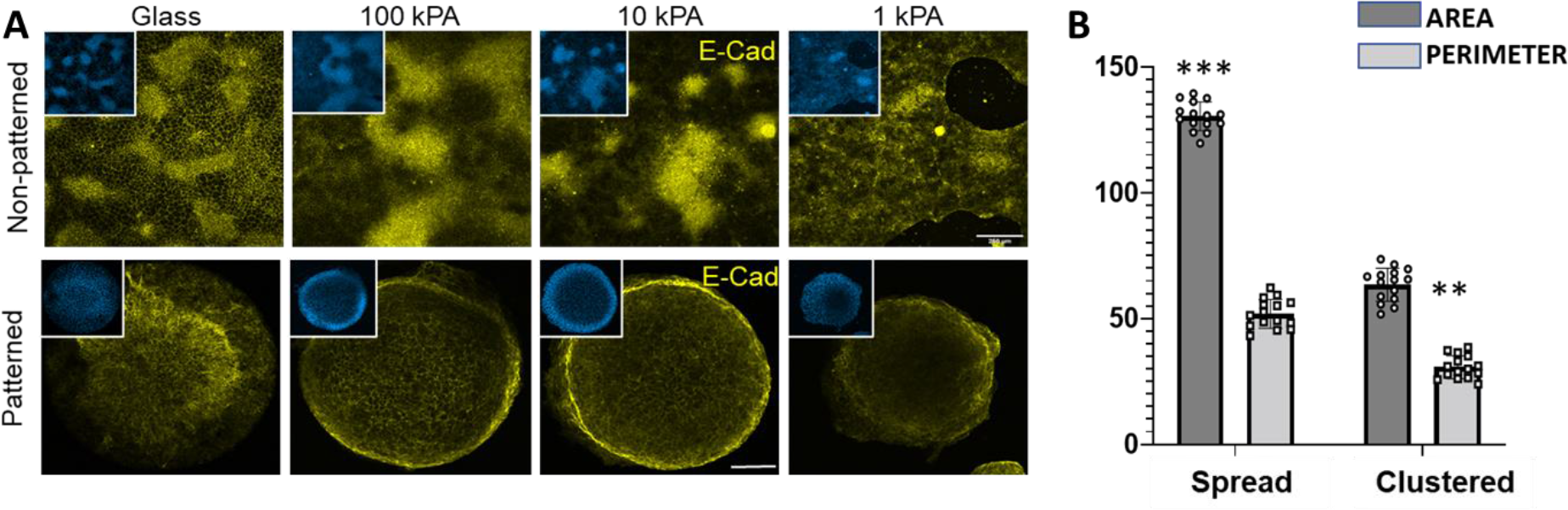
A) E-cad expression pattern on BMP4 treated non–patterned and patterned colonies. B) Nuclear size comparison for spread and clustered cells after 48hr treatment with BMP4 (N=15) Scale bars: 100µm

**Figure S10:**
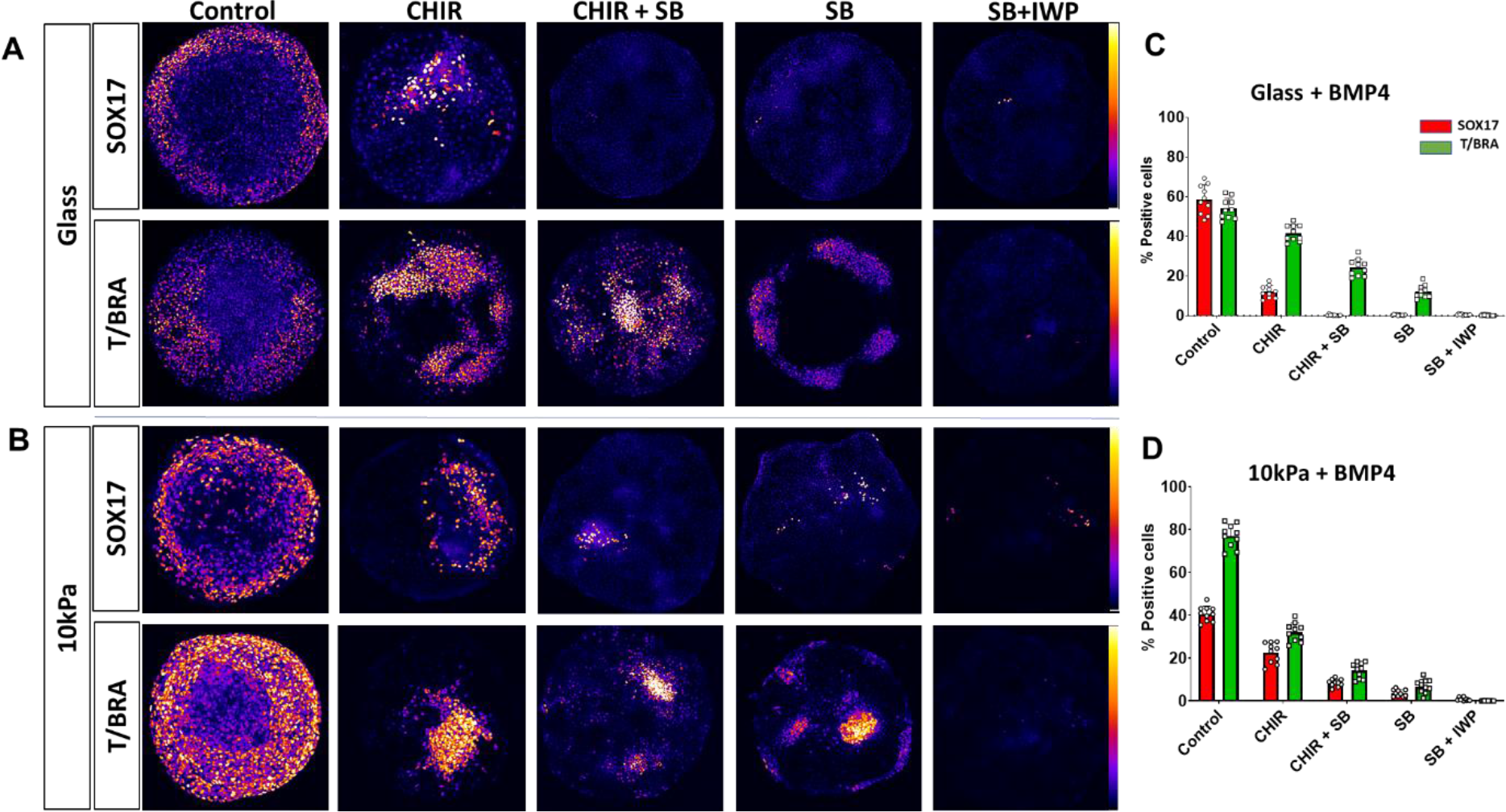
Small molecule inhibition on BMP4 sets: **A)** Heat maps for glass patterns at 48 hours – combinations of small molecule inhibitors for Wnt and Nodal SB = SB431542, IWP2 **B)** Heat maps for 10 kPa patterns at 48 hours – combinations of small molecule inhibitors for Wnt and Nodal **C)** Percent positive quantification for SOX17 and T/BRACHYURY in glass patterns + BMP4 N=10 p<0.001 **D)** Percent positive quantification for SOX17 and T/BRACHYURY in 10kPa patterns + BMP4 N=10 p<0.05 Scale bars - 100 µM.

**Figure S11:**
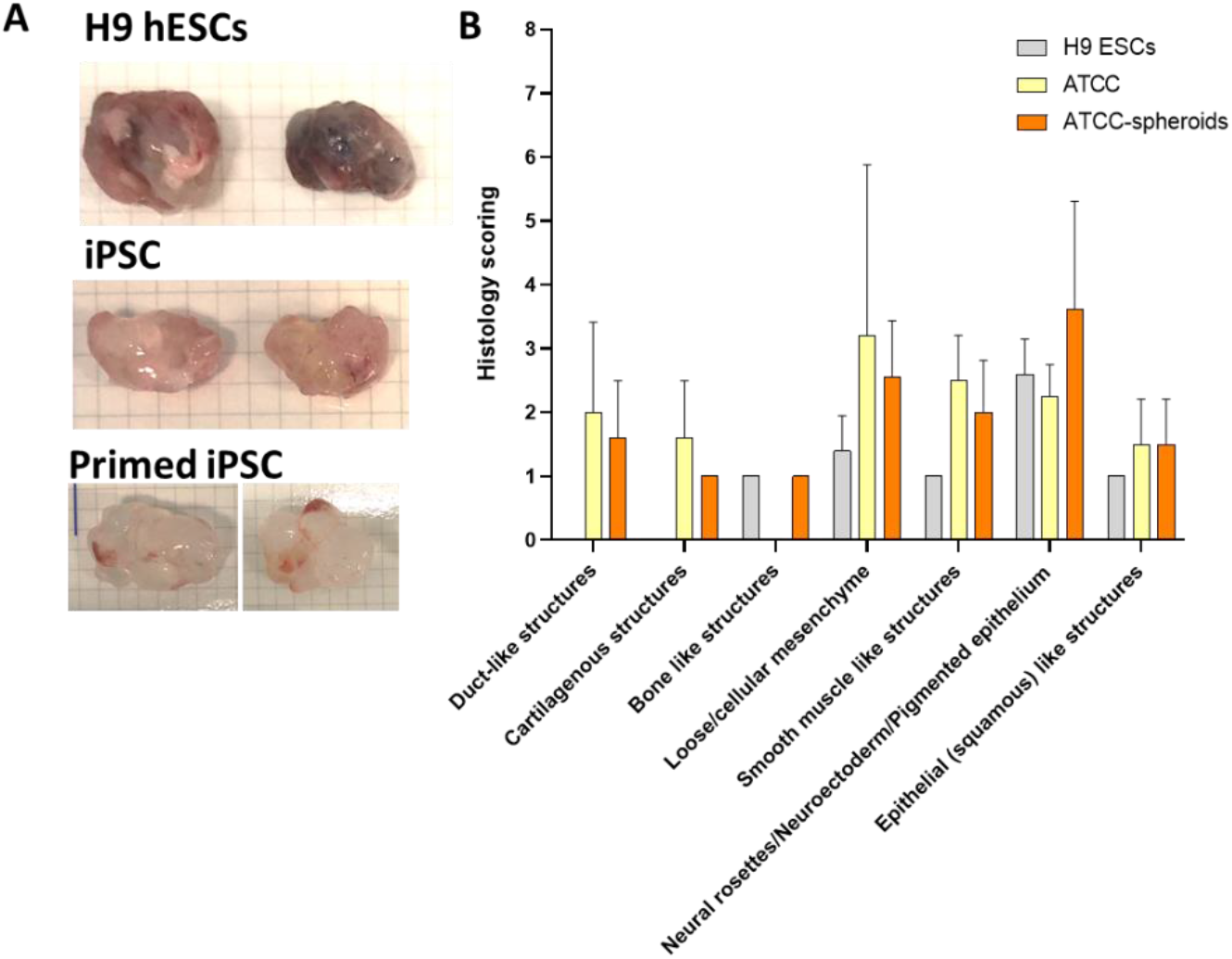
A) Comparative images for excised *in vivo* teratomas from H9 hESCs From ATCC hiPSCs from primed PA spheroids B) histology scoring of germ layer derivatives in all three groups

